# A new class of OLD family ATPases Cran1 functions in cell cycle progression in an archaeon

**DOI:** 10.1101/2025.03.27.645713

**Authors:** Yunfeng Yang, Shikuan Liang, Junfeng Liu, Xiaofei Fu, Pengju Wu, Haodun Li, Jinfeng Ni, Qunxin She, Mart Krupovic, Yulong Shen

## Abstract

Overcoming lysogenization defect (OLD) proteins are diverse ATPases-nucleases functioning in antiphage defense in bacteria. However, the role of these proteins in archaea is currently unknown. Here, we describe a new class of archaeal OLD family ATPases and show that they do not appear to be involved in antiviral defense but play an essential role in cell cycle progression. The gene in *Saccharolobus islandicus* REY15A, named as Cran1 (<u>C</u>ell cycle related <u>A</u>TPase and <u>n</u>ickase 1), cannot be deleted and exhibits cyclic expression patterns, with a peak expression during the transition from M-G1 to S cell cycle phase. Cran1 overexpression leads to significant growth retardation, cell size enlargement, and increased cellular DNA content. Cran1 displays potent nickase and ATPase activities *in vitro*, with the nickase activity being dependent on the presence of the ATPase domain. Notably, Cran1 copurifies with chromatin-associated proteins, such as Cren7 and a histone deacetylase homolog, suggesting its involvement in chromatin-related activities. Collectively our results suggest that Cran1 plays an important role in cell cycle progression, revealing a novel function of OLD family proteins.

## INTRODUCTION

Overcoming lysogenization defect (OLD) family proteins are ATP-powered nucleases that are widely distributed in bacteria, archaea and viruses (Dot *et al*, 2023). Recent bioinformatic, genetic, and structural studies have clarified the functioning and evolution of this protein family (Akritidou & Thurtle-Schmidt, 2023; Schiltz *et al*, 2020; Schiltz *et al*, 2019). These enzymes contain an N-terminal ATPase domain of the ATP-binding cassette (ABC) superfamily and a C-terminal TOPRIM (<u>Top</u>oisomerase-<u>prim</u>ase) domain nuclease (Aravind *et al*, 1998). OLD family proteins have been classified into four classes based on gene neighborhood, biochemical properties, domain organization, and catalytic mechanisms (Dot *et al*., 2023). Class 1 OLD family enzymes are encoded by standalone genes within variable gene neighborhoods and have been implicated in various virus-host and virus-virus antagonistic interactions. Indeed, the founding member of the family has been discovered in bacteriophage P2, with genetic studies showing that the *old*-bearing phage P2 kills *recB* and *recC* deficient *Escherichia coli* host strains upon infection and specifically interferes with phage λ replication (Lindahl *et al*, 1970). By contrast, class 2 OLD ATPases-nucleases are always encoded in tandem with UvrD/PcrA/Rep-like helicases (Doron *et al*, 2018), with the two enzymes, denoted GajA and GajB, respectively, constituting the recently described ‘Gabija’ antiviral defense system (Akritidou & Thurtle-Schmidt, 2023; Dot *et al*., 2023). Class 3 OLD ATPases-nucleases are commonly found in retrons, another defense system centered around a reverse transcriptase (*ret*) gene and a non-coding RNA (Gao *et al*, 2020; Mestre *et al*, 2020; Millman *et al*, 2020). Class 4 OLD enzymes constitute the PARIS defense system, with the ATPase and TOPRIM catalytic domains being encoded separately by adjacent genes, denoted *ariA* and *ariB* (Rousset *et al*, 2022). Although the properties of the OLD family enzymes have been explored extensively, focusing on the abortive infection of P2 lysogens originally discovered in the 1970s and, recently, on the action mechanisms of several defense systems (Antine *et al*, 2024; Cheng *et al*, 2023; Cheng *et al*, 2021; Doron *et al*., 2018; Schiltz *et al*., 2020; Schiltz *et al*., 2019; Yang *et al*, 2024), it remains unclear whether the OLD family enzymes have any non-antiviral functions *in vivo*. To the best of our knowledge, none of the archaeal OLD ATPases-nucleases has been characterized and reported.

Archaea can be divided into four supergroups or superphyla, namely, Euryarchaeota, TACK superphylum, Asgardarchaeota, and DPANN superphylum (Liu *et al*, 2021c; Spang *et al*, 2017). In addition to sharing common origin of information processing machineries with Eukarya, archaea of the Asgardarchaeota and TACK superphylum encode many proteins that were initially thought to be specific of eukaryotes (Imachi *et al*, 2020; Liu *et al*., 2021c). In particular, species of the order Sulfolobales from the TACK superphylum have a cell cycle resembling that of eukaryotes (Gomez-Raya-Vilanova *et al*, 2025; Lundgren & Bernander, 2005; Lundgren *et al*, 2008), including G1 (Gap 1), S (DNA synthesis), G2 (Gap 2), M (chromosome segregation), and D (cell division) phases (Lindås & Bernander, 2013). A newborn Sulfolobales cell undergoes a brief interphase G1 (about 5% of the entire cell cycle) during which the cell grows and prepares for genome replication. The cell then enters the S phase, which accounts for about 30% of the cell cycle, during which the genome is duplicated. Then, the cell undergoes a long G2 phase (about 60%), during which cells are actively metabolizing and increase in size, preparing for genome segregation (M phase) and cell division (D phase), which follow each other in quick succession.

The cell division machinery of Sulfolobales consists of archaea-specific protein CdvA, four eukaryotic ESCRT-III homologs, namely, ESCRT-III (also known as CdvB), ESCRT-III-1 (CdvB1), ESCRT-III-2 (CdvB2), and ESCRT-III-3 (CdvB3), and the AAA^+^ ATPase Vps4 homolog (CdvC) (Lindås *et al*, 2008; Liu *et al*, 2017; Risa *et al*, 2020; Samson *et al*, 2008). Although the role of ESCRT machinery in the Sulfolobales cytokinesis has been firmly established, the regulation of the cell cycle progression remains unclear. The expression of the *cdvA* is transcriptionally controlled by a repressor aCcr1 (Li *et al*, 2023; Yang *et al*, 2023b), whereas the subsequent degradation of CdvB by the proteasome is reported to control the cytokinesis progression (Liu *et al*, 2025a; Risa *et al*., 2020). However, how the other cell cycle phases, especially G1 and S, are regulated remains unknown. Interestingly, in contrast to Sufolobales cell, eukaryotic cells have a relatively long G1 phase, during which the activation of different cyclins and Cdk proteins determines whether the cell will enter the cell cycle or remain in a quiescent state (Matthews *et al*, 2022). Because eukaryotes are believed to have originated from archaea, elucidating the cell cycle regulation in archaea, including the regulation of the M-G1 and S phases, could provide insights into the origin and evolution of the eukaryotic cell cycle regulation mechanisms.

In this study, we characterize an OLD family protein from *S. islandicus* REY15A (order Sulfolobales), one of the few genetically tractable representatives of the TACK superphylum. We show that the gene coding for Cran1 is conserved across Sulfolobales and could not be deleted, suggesting its essentiality for the cell viability. Unexpectedly, Cran1 does not appear to participate in antivirus defense, but instead is involved in cell cycle progression. Cran1 exhibits cyclic expression patterns at both transcriptional and translational levels, and its overexpression impairs cell cycle progression. Interestingly, the ATPase activity, but not the nuclease activity appear to be critical for this function. Our study expands the understanding on the functional versatility of the OLD family proteins and provides insight into the cell cycle regulation in Sulfolobales.

## Methods

### Strains and growth conditions

*Saccharolobus islandicus* E233S(Δ*pyrEF*Δ*lacS*) (Reagents and tools table) was grown in STVU medium containing mineral salt, 0.2% (w/v) sucrose (S), 0.2% (w/v) tryptone (T), 0.01% (w/v) uracil (U) and a mixed vitamin solution (V). The medium was adjusted to pH 3.3 with sulfuric acid, as described previously (Deng *et al*, 2009). STV medium containing 0.2% (w/v) tryptone (T) was used for screening and cultivating uracil prototrophic transformants. ATV medium containing 0.2% (w/v) D-arabinose(A) was used for protein expression. Culture plates were prepared using gelrite (0.8% [w/v]) by mixing 1 × STV and an equal volume of 1.6% gelrite. Plasmid cloning and protein expression were performed with *E. coli* DH5α and BL21 (DE3)-RIL, respectively. The cells were cultured in Lysogeny-Broth (LB) medium supplemented with appropriate antibiotics. Plaque assay was performed using *E. coli* MG1655. The strains constructed and used in this study are listed in Appendix Table S1.

### Phylogenetic analysis

The phylogenetic analyses were similar to those previously described (Yang *et al*., 2023b). In short, a dataset of OLD family proteins from bacteria and archaea were collected by PSI-BLAST (2 iterations against the RefSeq database at NCBI; E =1e-05)(Altschul *et al*, 1997)(Appendix Table S3). The collected sequences were then clustered using MMseq2 to 90% identity over 80% of the protein length. In the case of Class 4 OLD proteins, the ATPase and nuclease subunits were concatenated. Sequences were aligned using MAFFT v7 (Katoh *et al*, 2019) and the resultant alignment trimmed using Trimal (Capella-Gutiérrez *et al*, 2009), with the gap threshold of 0.2. Maximum likelihood phylogenetic analysis was performed using IQ-Tree (Minh *et al*, 2020), with the best selected amino acid substitution model being LG + I + G4. The branch support was assessed using SH-aLRT (Guindon *et al*, 2010).

### Construction of plasmids

Construction of *cran1* knockout and *in situ* gene tagging of *cran1* strains was according to the endogenous CRISPR-Cas-based genome editing method as described previously (Li *et al*, 2016). A target site was selected on the *cran1* consisting of a 5’-CCN-3’ type I-A protospacer neighboring motif and a 40 nt downstream DNA sequence (protospacer). Two complementary DNA oligonucleotides were then designed based on the protospacer and synthesized. The spacer fragments were generated by annealing the corresponding complementary oligonucleotides and inserted into the BspQI site of pGE, producing plasmids carrying the artificial mini-CRISPR arrays. The donor DNA fragment was generated by splicing and overlapping extension (SOE)-PCR with *ApexHF* HS DNA Polymerase FS (Accurate Biotechnology, Hunan, China). The Donor DNA fragments were ligated to the XhoI and SphI cleavage sites of pGE by the TEDA method(Xia *et al*, 2019) generating the editing plasmid.

Cran1 and its mutants were overexpressed in *Sa. islandicus* using the pSeSD vector (Peng *et al*, 2012; Peng *et al*, 2017) and in *E. coli* using the pET22b vector. PCR was used to amplify the target fragment and overlapping extension PCR was used to obtain the site-specific mutants. The *cran1* knockdown vector was constructed by using tandem repeats having multiple spacers. The spacer sites were chosen to contain 40 bp (TAGATTAGTTTTTCCATATCCGTTATAGCCAACGACTACG) immediately after the antisense strand GAAAG (Peng *et al*., 2017) and the multiple spacers were separated by the repeat sequences. The knockdown plasmid was transferred into *in situ* gene tagging of *cran1* strain for the detection of Cran1 levels with anti-Flag antibody. The primers used in this study are listed in the Appendix (Appendix Table S2).

### Bright-field microscopy

For bright-field microscopy analysis, 5 μl of cell suspension at the indicated time points were examined under a NIKON TI-E inverted fluorescence microscope (Nikon, Japan) in Differential interference contrast (DIC) mode.

### Flow cytometry analysis

The cell cycle of synchronized cells of E233S carrying pSeSD and the Cran1 overexpression plasmid was analyzed by flow cytometry using an ImageStreamX MarkII quantitative imaging analysis flow cytometer (Merk Millipore, Germany). Briefly, cells were fixed with ethanol at a final concentration of 70% for at least 12 h at each corresponding time point and stained using propidium iodide (PI) at a final concentration of 50 μg/ml. The data were collected for at least 20,000 cells per sample and analyzed using IDEAS data analysis software.

### Cell cycle synchronization

In situ gene tagging of *cran1* strains were synchronized as previously described (Liu *et al*, 2021b; Yang *et al*., 2023b). Briefly, cells were first grown aerobically at 75□ with shaking (145 rpm) in 30 ml of STVU medium. When the OD_600_ reached 0.6-0.8, the cells were transferred into 100 ml STVU medium with an initial estimated OD_600_ of 0.05 and cultivated as above. When the OD_600_ reached 0.15-0.2, acetic acid was added at a final concentration of 6 mM and the cells were blocked at G2 phase of the cell cycle after 6 h treatment. Then, the cells were collected by centrifugation at 3000 g for 10 min at room temperature to remove the acetic acid and washed twice with 0.7% (w/v) sucrose. Finally, the cells were resuspended into 100 ml of pre-warmed STVU medium and cultivated as above for subsequent analysis. For the synchronization of cells carrying plasmid for Cran1 and mutant overexpression, the ATV medium was used to induce protein expression after removal of acetic acid. The cell cycle was analyzed using flow cytometry.

### Protein expression and purification

Recombinant Cran1 and mutants with C-terminal His-tag were expressed using *Escherichia coli* BL21 (DE3)-RIL cells. Protein expression was induced during logarithmic growth (OD_600_=0.4∼0.8) by the addition of 0.5 mM isopropyl-thiogalactopyranoside (IPTG) followed by an overnight cultivation at 16□. Similarly, proteins were purified from *Sa. islandicus* using Cran1 and its mutant overexpression strains. When the overexpression strains were grown in MTV medium to OD_600_=0.2, the expression was induced by adding arabinose and the cultivation continued for 24h. Cells were collected by centrifugation and the cell pellet was resuspended in buffer A (50 mM Tris-HCl pH 8.0, 300 mM NaCl, 5% Glycerol). Cells were lysed by sonication and the cell extract was clarified by centrifugation at 13 000 × g for 20 min at 4□. The supernatant was incubated at 70□ for 20 min (for expression in *E. coli* only), centrifuged at 12, 000 g for 20 min again, and then filtered through a membrane filter (0.45 μm). The samples were loaded on to a Ni-NTA agarose column (Invitrogen) pre-equilibrated with buffer A and eluted with a linear imidazole (40-300 mM imidazole) gradient in buffer A. Fractions were pooled and concentrated to 1 ml and purified further by size exclusion chromatography (SEC) using a Superdex 200 column using the buffer A (50 mM Tris-HCl pH 8.0, 300 mM NaCl, 5% Glycerol). Finally, the samples were dialyzed in a storage buffer containing 50 mM Tris-HCl pH 7.4, 100 mM NaCl, 1mM DTT, 0.1 mM EDTA and 50% glycerol. Protein concentrations were determined using the Bradford protein assay kit (Beyotime) and the purity was analyzed by 15% SDS-PAGE stained with Coomassie blue.

### Western blotting

The expression levels of Cran1 and cell division proteins in the synchronized cells were analyzed by Western blotting. Around 2×10^8^ cells were collected at the indicated time points for each sample and subjected to SDS-PAGE analysis in a 15% gel. The separrated proteins were then transferred onto a PVDF membrane. The specific bands were detected by chemoluminescence using Sparkjade ECL super Western blotting detection reagents (Shandong Sparkjade Biotechnology Co., Ltd.) according to the manufacturer’s instructions. The primary antibodies against ESCRT-III, ESCRT-III-1, and TBP were produced in rabbit (HuaAn Biotechnology Co., Hangzhou, Zhejiang, China)(Liu *et al*., 2021b; Liu *et al*., 2017), and the primary antibody against Flag tag and His tag were purchased from TransGen Biotech company (Beijing, China). The goat anti-rabbit antibodies (KermeyBiotech,Zhengzhou,China) coupled with peroxidase were used as secondary antibodies.

### Electrophoretic mobility shift assay (EMSA)

The 100 nt FAM-labelled Cran1 promoter sequence (Appendix Table S2) were used as the substrates to determine the binding capacity of aCcr1. The binding assay was performed in a 20 μl reaction mixture containing 10 nM dsDNA, 50 mM Tris-HCl, pH 7.4, 5 mM MgCl_2_,20 mM NaCl, 50 μg/mL BSA, 1 mM DTT, 5% glycerol, and different concentrations of purified proteins. The reaction mixture was incubated at 37 □ for 30 min before loaded onto a 12% native polyacrylamide gel. After running in 0.5×TBE, the gel was visualized using an Amersham ImageQuant 800 biomolecular imager (Cytiva).

### DNA cleavage assays

Supercoiled pUC19 plasmid (300 ng) was with 2 μM protein in a final volume of 20 µl in a DNA cleavage buffer (20 mM Tris-HCl pH 7.0, 50 mM KCl, 0.1 mg/ml BSA, and 10 mM divalent metal chloride salt). Reactions were carried out at 70□ for 30 min and then the samples were quenched with 5 μl of 0.5 M EDTA, pH8.0. Samples were analyzed via 1% native agarose gel electrophoresis. Supercoiled and nicked plasmids were extracted using the OMEGA Plasmid Extraction Kit, and linearised plasmids were generated by HindIII single restriction. The signal of the initial DNA substrate (linearized or supercoiled plasmid) was measured and quantified using the ImageJ software if required. The site-directed nuclease and ATPase mutants were assayed under optimal divalent metal conditions of 10 mM MgCl_2_ and 1 mM CaCl_2_. The cleavage efficiency was quantified by comparing the band intensity in each lane and calculating the percentage of DNA digested relative to the control.

### ATPase assays

The ATPase activity was examined using a colorimetric malachite green assay that monitored the amount of free phosphate released over time (Monroe *et al*, 2014). The reactions in a mixture containing 0.2 μM Cran1 in 25 mM Tris-HCl pH 7.4, 100 mM NaCl, 5 mM MgCl_2_, and 1 mM ATP in a total volume of 50 μl were carried out for at least 5 min at indicated temperatures (30, 40, 50, 60, 70, 80, or 90□). The reaction mixture was then placed on ice and quenched with 100 μl of malachite green color reagent (14 mM ammonium molybdate, 1.3 M HCl, 0.1% (v/v) Triton X-100, and 1.5 mM malachite green) and 50 μl of 21 % (w/v) citric acid. The green compound formed by malachite green, molybdate, and free phosphate was detected by absorbance at 650 nm using spectrophotometer. A sodium phosphate standard curve was used to estimate the amount of phosphate released during ATP hydrolysis. NTP hydrolysis activity including the ATPase activity assays of Cran1 site mutants were performed at 70□. It should be noted that above 70□, ATP is partially hydrolyzed by heat, so the amount hydrolyzed by heat were excluded from the calculation of ATP hydrolysis activity at 70, 80, and 90□.

### Spot assays

For the phage infection assay of *E. coli*, *cran1* sequence or *cran1* with *sire_0087* were cloned into the pET22b generating a vector for the expression of recombinant Cran1. The plasmid was transformed into *E. coli* MG1655. A colony was picked up from the plate and grown in LB broth containing ampicillin (100 mg/ml) at 37 □ to an OD_600_ of 0.4. Protein expression was induced by the addition of 0.2 mM IPTG. After 1 hour of growth, 500 µl of bacterial culture was mixed with 10 ml of 0.5% LB top layer agar and the whole sample was poured onto an LB plate containing ampicillin (100 mg/ml) and IPTG (0.1 mM). Plates were spotted with 3 µl of six 10-fold dilutions of phage T1-T7 of 10^0^-10^-5^ diluted in LB liquid medium. The plates were incubated overnight at 37□ and then imaged. The spot assay of *Sa. islandicus* REY15A was according to previous report (Liu *et al*, 2021a). Briefly, the control cells (Sis/pSeSD), Cran1 expression, and the Cran1 and sire_0087 co-expression cells were collected at mid-logarithmic phase and mixed with 10 ml of pre-heated ATYS medium containing 0.4% (wt/vol) phytagel, vortexed and immediately poured into the empty Petri dishes. After the plates solidified, 10µl of the 10-folds serially diluted STSV2 or SMV1 preparation were applied on the plates and incubated at 75 □ for 2-3 days.

### Chromatin fractionation

Chromatin fractionation analyses were similar to those previously described (Takemata *et al*, 2019). In brief, the cells were harvested from the cultures at OD_600_ = 0.4 for exponential samples and OD_600_ = 1.0 for stationary samples. The cells were resuspended in 100 μl/1 OD_600_ unit/ml of chromatin extraction buffer (25 mM HEPES, 15 mM MgCl_2_, 100 mM NaCl, 400 mM sorbitol, 0.5% Triton X-100, pH 7.5) and incubated on ice for 10 minutes. The extracts were centrifuged for 20 minutes at 14,000 *g*, 4 □. The soluble fraction was transferred to a new tube and the pelleted chromatin fraction was resuspended in a volume of chromatin extraction buffer equivalent to the soluble fraction. The chromatin fraction was then sonicated three times for 30 s each.

### Immunoprecipitation and mass spectrometry

1L of *Sa. islandicus* culture (Cran1 *in situ* Flag-tagged strain) in logarithmic growth phase (OD_600_=0.3∼0.6) was collected. The cells were suspended in 1 mL of lysis buffer (50 mTris-HCl, pH6.8, 150 mM NaCl, 0.1 mM DTT, 10%, w/v, glycerol, 0.5%) containing protease inhibitor PMSF (Sigma-Aldrich). The cells were broken by ultrasonication. The sample was centrifugated at 20,000×*g* at 4°C for 30 min and the supernatant was collected and used as the crude extract (input). Then RNase A was added into 500 μL of crude cell extract. The sample incubated on ice for 30 min to remove RNA. Next 50 μL anti-FLAG® M2 magnetic beads (Sigma-Aldrich) were added to the sample and the tube was incubated at 4°C with shaking for 1 hr. The beads were collected by using the magnetic rack and the supernatant (FT, flow through) was discarded. The magnetic beads were washed 5 times with lysis buffer. Finally, the proteins were eluted with 100 μl of 0.1 M glycine HCl, pH 3.0. The eluate was added into 10 μl Tris (1M), pH 8.0 to protect the eluted proteins from prolonged periods of time in the pH 3.0 solution. For The eluate sample was added into 20 μL of 5× loading buffer (100 mM Tris-HC1, pH 6.8, 200 mM β-mercaptoethanol, 4% SDS, 0.2% bromophenol blue, 20% glycerol). After boiled for 10 min, the tube was centrifugated and the supernatant was subjected for SDS-PAGE. The bands were sliced and subjected for LC-MS analysis at Beijing Protein Innovation Co., Ltd. The identified proteins are listed in Appendix Table S4.

## RESULTS

### Phylogenetic diversity of OLD family proteins

All currently functionally characterized OLD family enzymes, assigned to four classes, are encoded by bacteria or bacterial viruses and are implicated in antiviral defense (Dot *et al*., 2023). To assess the diversity and taxonomic spread of this protein family, we assembled a dataset of OLD family proteins from bacteria and archaea (Appendix Table S3) and performed maximum likelihood phylogenetic analysis. Notably, not all four classes of characterized OLD family proteins were monophyletic. In particular, Class 2 enzymes were nested within the diversity of Class 1 proteins, whereas Class 4 proteins in which the ATPase and nuclease domains are split, were nested within Class 3, as a sister group to subgroup 3b. These observations are consistent with the notion that new defense systems evolve through shuffling of different modules through recombination (Koonin *et al*, 2017). Whereas Classes 1 and 2 were distributed in both prokaryotic domains, Classes 3 and 4 were restricted to bacteria (Appendix Table S3). Notably, phylogenetic analysis uncovered at least 11 well-supported, previously uncharacterized clades (U1-U11) of OLD family proteins (Fig. 1A). Three of these clades (U1, U2 and U4) included both bacterial and archaeal representatives, five clades (U5, U6, U7, U9 and U10) were dominated by diverse bacterial sequences and three clades (U3, U8 and U11) had nearly exclusively archaeal membership (Appendix Table S3). Mixed phyletic patterns (e.g., mixed bacterial and archaeal membership) observed in some of the clades are consistent with the horizontal spread characteristic of defense systems (Rocha & Bikard, 2022). By contrast, clades U8 and U11 were homogeneous in their taxonomic representation, restricted to hyperhalophilic (order Halobacteriales) and thermoacidophilic (order Sulfolobales) archaea, respectively. Notably, the phylogenetic relationship between the OLD family sequences in U11 fully recapitulated the species tree of Sulfolobales (Fig. 1B) (Counts *et al*, 2021), suggesting conservation and vertical inheritance of the corresponding genes in this archaeal order, without obvious signs of horizontal gene transfer. Intrigued by such atypical conservation of the potential OLD family defense genes in Sulfolobales, we set out to characterize them functionally.

**Figure 1.**
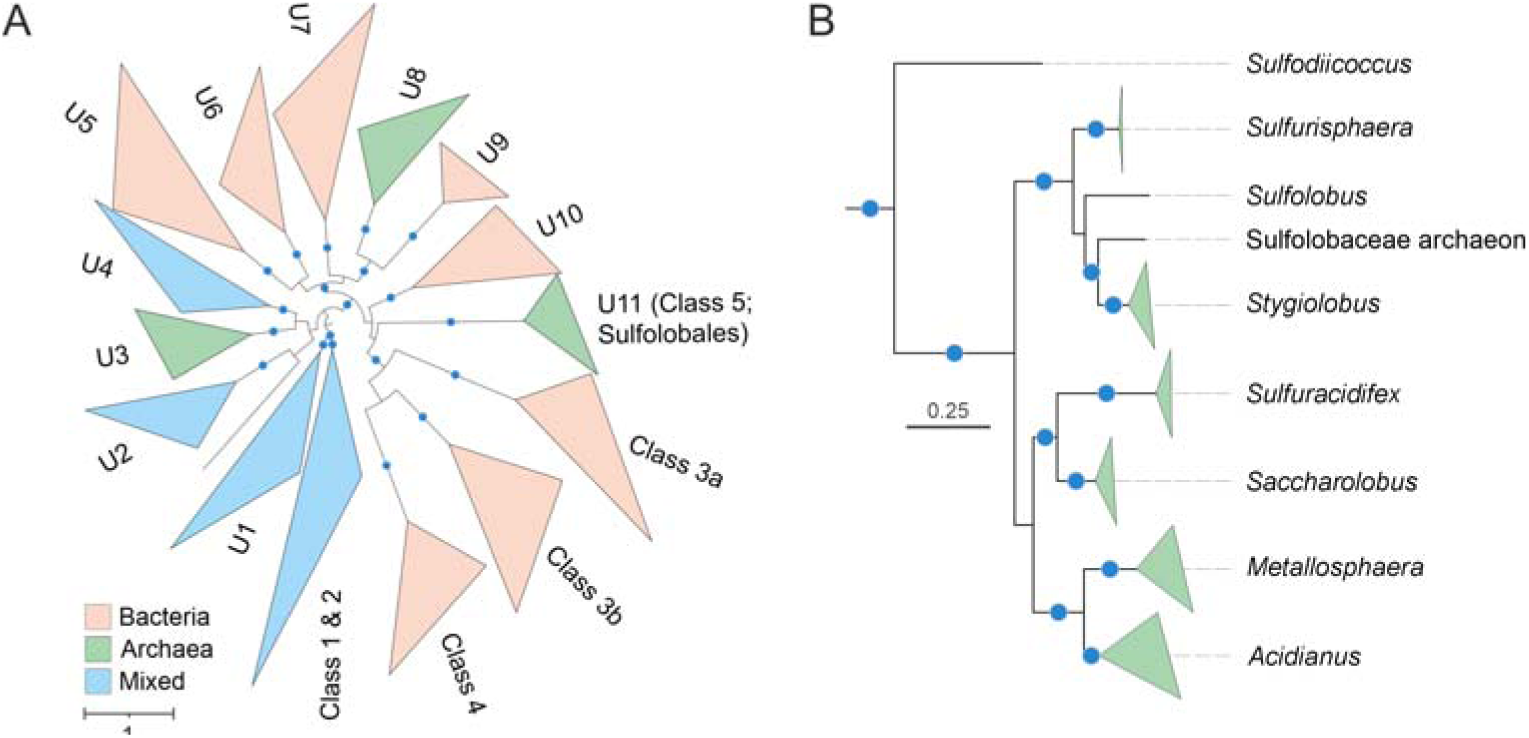
Diversity of OLD family proteins in bacteria and archaea. (**A**) Maximum likelihood phylogenetic analysis of OLD family proteins. The tree was midpoint rooted. Previously characterized groups of OLD family proteins are denoted by Classes 1 through 4, whereas the clades encompassing uncharacterized protein sequences are labeled U1-U11. Clades which included >90% of proteins encoded by organisms of the same domain were considered as bacterial or archaeal, respectively; when the clades included >10% of proteins encoded by organisms from the other domain, they were denoted as mixed. (**B**) Detailed view of the U11 clade including OLD family proteins conserved in Sulfolobales. In both panels, well supported clades were collapsed to triangles, whose side lengths are proportional to the distances between closest and farthest leaf nodes. The scale bars represent the number of substitutions per site, whereas blue circles denote SH-aLRT (Shimodaira–Hasegawa approximate likelihood ratio test) branch supports higher than 90.

### Sulfolobales OLD proteins have expected domain organization and are catalytically active

Analysis of a representative structural model of the Sulfolobales OLD proteins revealed a domain organization typical of this protein family, with the N-terminal ABC family ATPase and the C-terminal TOPRIM nuclease domains. Akin to most other OLD family ATPases, with a notable exception of Class 1 enzymes, Sulfolobales OLD proteins have an α-helical insertion within the TOPRIM domain (Figure 2A and 2B). Multiple sequence alignment of the Sulfolobales OLD homologs has shown that all active site residues in both ATPase and nuclease domains are conserved (Appendix Figure S1), suggesting that the protein is enzymatically active.

**Figure 2.**
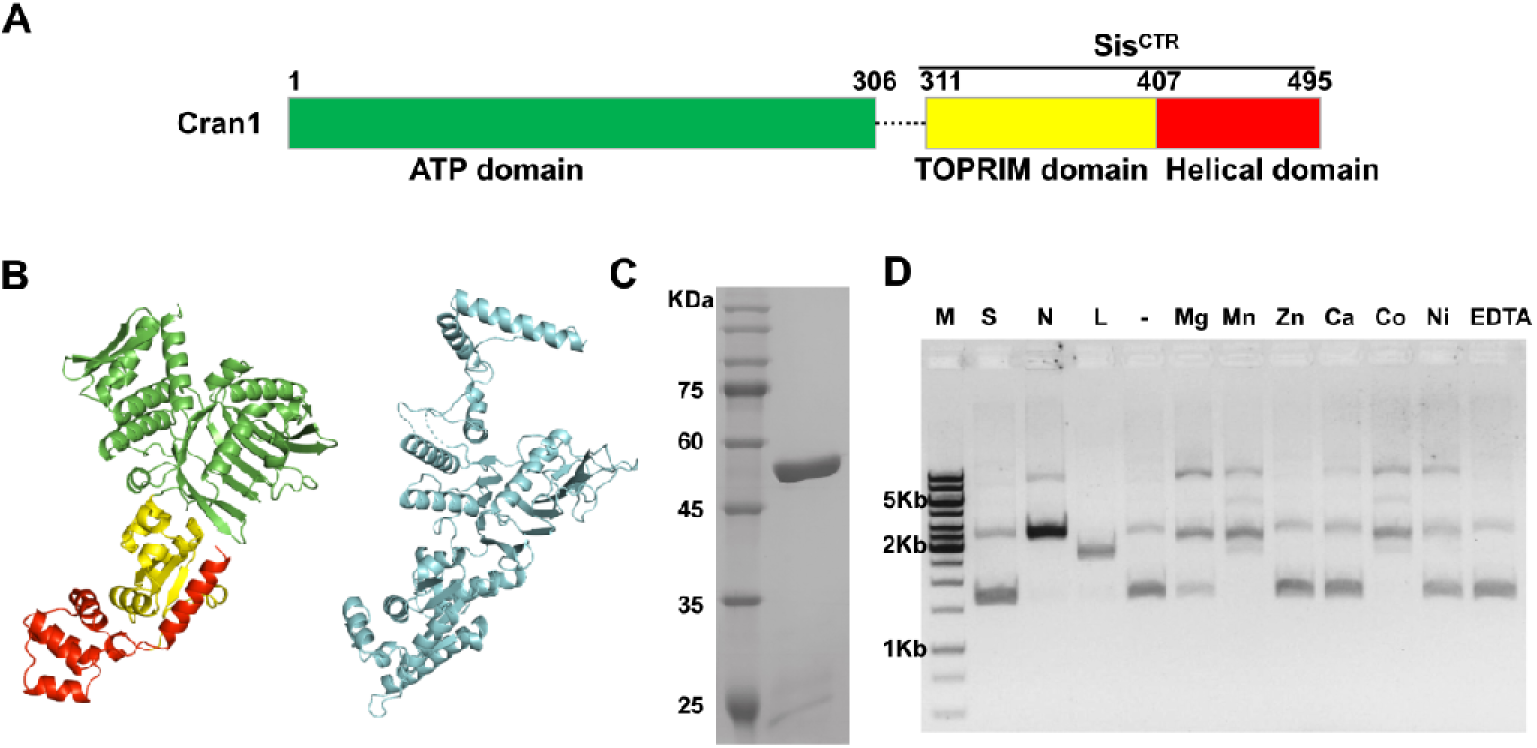
Cran1 exhibits metal-dependent nuclease activity *in vitro*. (**A**) Domain architecture of Cran1. Sis^CTR^, the C-terminal domain of Cran1. (**B**) Structural model of Cran1 predicted by Alphafold3 (ipTM = 0.96), with different colours representing different structural domains as in (A) and the Class 1 OLD family protein from *Thermus scotoductus* (blue, PDB:6p74_A). (**C**) SDS-PAGE showing purified full-length Cran1. (**D**) Metal-dependent nuclease activity of Cran1 on supercoiled pUC19. ‘N’, ‘L’, and ‘S’ denote the positions of ‘nicked’, ‘linearized’, and ‘supercoiled’ pUC19 plasmid, respectively.‘M’, DNA marker.‘-’, no divalent metal ions added. ‘EDTA’, EDTA replacing divalent metal ions. For more detailed experimental procedures, see the Methods section.

To test the *in vitro* activities of Sulfolobales OLD experimentally, we focused on an OLD protein SiRe_0086 (Cran1) encoded by *Saccharolobus islandicus* REY15A, a thermoacidophilic archaeon growing optimally at 76-80□ and pH of 2-3. The Cran1 protein was expressed and purified from *Sa. islandicus* cells (Figure 2C). Notably, whereas OLD family ATPases typically form dimers, in the gel filtration chromatography analysis, Cran1 eluted at a peak of 15.1 ml (Figure EV1A), corresponding to a monomeric form (55 kDa).

*Nuclease activity*. The TOPRIM domain nucleases usually require divalent metal ions for catalysis (Yang, 2011). Thus, we examined the nuclease activity using supercoiled and linearized forms of plasmid pUC19 as the substrates in a buffer containing different divalent metal ions (Figure 2D). In the presence of 10 mM Mn^2+^ and Co^2+^, Cran1 exhibited strong nickase activity, cleaving nearly 100% of the substrate within 20 min (Figure 2D). In the presence Mg^2+^, ∼70% of the supercoiled pUC19 plasmid substrate was nicked within 20 min. By contrast, very week activity was detected in the presence of Ni^2+^ and no activity was observed with Ca^2+^ and Zn^2+^ as the cofactor (Figure 2D). These data indicate that specific divalent cations are required for Cran1 activity. Notably, Ca^2+^ alone does not promote nuclease activity on supercoiled DNA substrates; however, with a combination of Ca^2+^ and Mg^2+^, the activity of Cran1 was higher than with either of these ions alone (Figure EV1B). The enzyme showed the strongest nuclease activity in the presence of 10 mM Mg^2+^ and 1 mM Ca^2+^. Therefore, this combination of divalent cations was used in the subsequent experiments.

Cran1 showed strong nickase activity at a broad range of temperatures (between 40□ and 70□; Figure EV1C) and pH (pH 5-10, optimum at pH 6-8; Figure EV1D). Notably, neither ATP nor its derivatives (ADP, dATP, and AMP) had a measurable effect on the nuclease activity of Cran1 when added to the reaction mixtures (Figure EV1E). However, the mutants in which the ATPase domain was deleted or magnesium ion binding/ATP hydrosis site was mutated lost the nuclease activity (see below), suggesting that active ATPase domain is necessary for Cran1 to function as a nuclease. Notably, the nuclease activity of Cran1 was inhibited by high salt concentrations (Figure EV1F).

Our results indicate that Cran1 has similar enzymatic properties as the reported Class 2 OLD family proteins. Based on the previous reports on the mutagenesis of a Class 2 OLD family nuclease (Schiltz *et al*., 2019) and our structural modelling, we hypothesized that Cran1 uses a two-metal ion catalytic mechanism and predicted that E338, L391, D393, and E341, E409, and E411 are responsible for the binding of metal ions A and B (Figure 3A), respectively. To test this hypothesis, we constructed three mutants, 3A (E338A/L391A/D393A), 3B (E341A/E409A/E411A), 2A2B (L391A/D393A/E409A/E411A), and purified the corresponding proteins (Figure EV2A). The three mutants completely lost the nuclease activity on pUC19 plasmid (Figure 3B), suggesting that both metal binding sites are essential for the nuclease activity.

**Figure 3.**
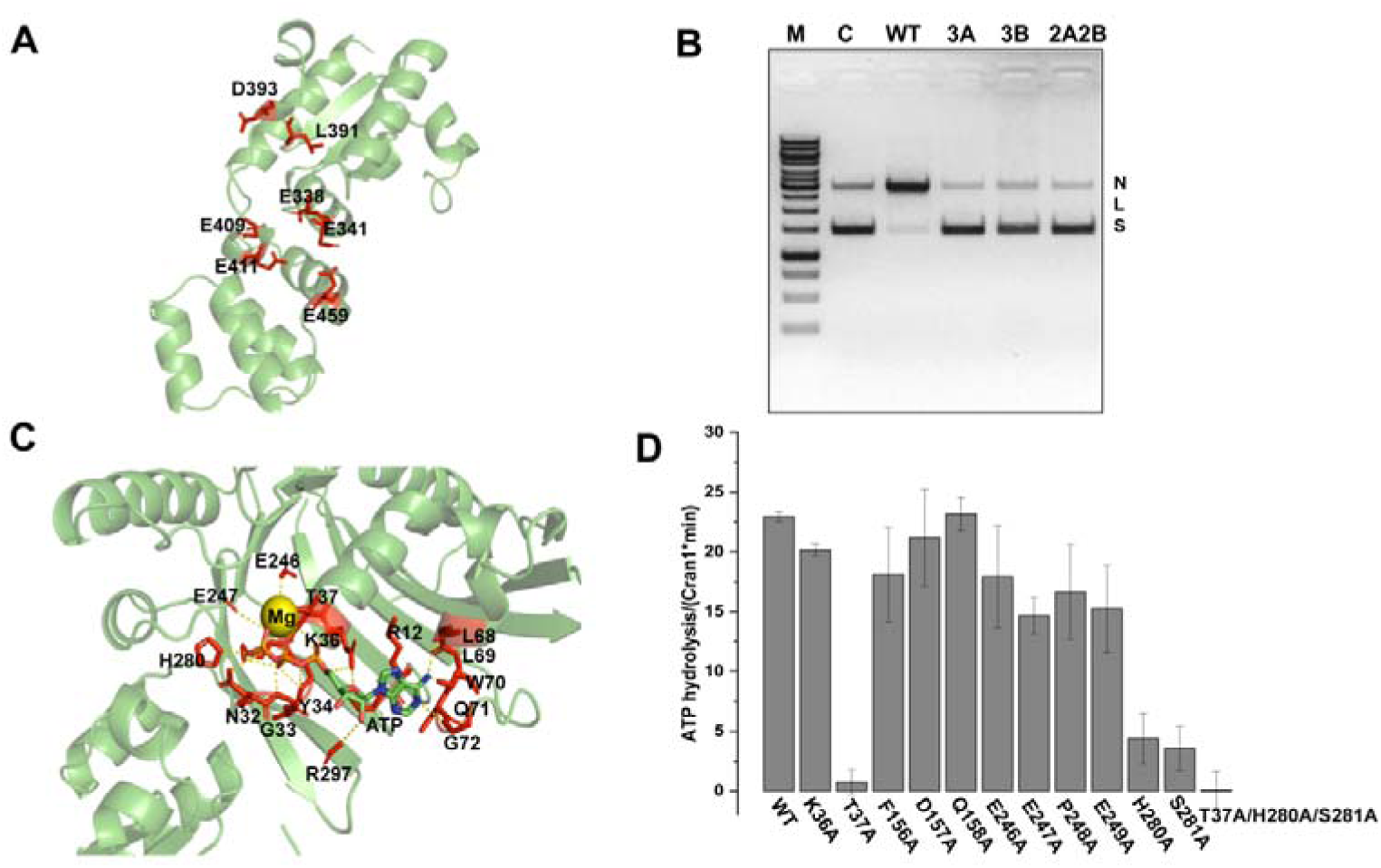
Cran1 nuclease active site and ATPase key site mutants. (**A**) Metal ion binding residues in the structural domains of nucleases predicted based on protein structure and sequence. (**B**) Analysis of nickase activity of key residue combination mutants 3A, 3B, 2A2B of nuclease *in vitro*. WT represents wild-type Cran1 protein. 3A represents the E338A, L391A, D393A combination mutant, 3B represents the E341A, E409A, E411A combination mutant, and 2A2B represents the L391A, D393A, E409A, E411A combination mutant. ‘N’, ‘L’, and ‘S’ denote the positions of ‘nicked’, ‘linearized’, and ‘supercoiled’ DNA, respectively. (**C**) Structure showing the key amino acid residues (highlighted in red with atomic distances within 5 Å) in the active center of the Cran1 ATPase domain predicted by AlphaFold3. (**D**) Analysis of ATP hydrolytic activity of the mutants of key amino acids of ATPase *in vitro*.

*ATPase activity*. Although all OLD family proteins have a typical ATPase domain, not all OLD proteins have ATPase activity (e.g., GajA) (Cheng *et al*., 2021). To determine whether Cran1 is capable of hydrolyzing ATP, we expressed and purified Cran1 from *Sa. islandicus* and examined its ATPase activity *in vitro*. As shown in Figure EV2C and D, Cran1 has ATPase activity with an optimal activity at 70-80□. We identified the key amino acid residues for the ATPase activity by analyzing the structural model generated using AlphaFold3 and sequence alignment (Figure 3C and Appendix Figure S1). The proteins in which the conserved residues were individually mutated to alanine were expressed and purified (Figure EV2B), and their ability to hydrolyze ATP was examined (Figure 3D). The ATPase activity of T37A, H280A, and S281A mutants was severely impaired, with T37A being the least active, whereas the triple mutant T37A/H280A/S281A completely lost the ATPase activity (Figure 3D). Notably, the latter mutant also lost the nuclease activity, suggesting a crosstalk between the ATPase and nuclease domains. Consistently, deletion of the ATPase domain rendered the nuclease domain inactive (Figure EV2E). Collectively, these results indicate that Cran1 is a functional representative of the OLD family, endowed with both nuclease and ATPase activities. Thus, we denote the clade U11 conserved across Sulfolobales as class 5 (Figure 1).

### Sulfolobales OLD family proteins do not appear to be involved in antivirus defense

Bacterial class 2-4 OLD enzymes are encoded within conserved genomic loci including other components of the corresponding defense systems. Genomic neighborhood analysis of the Sulfolobales *old* genes shows that they form a conserved potential operon with a gene encoding a predicted membrane protein of unknown function (*sire_0087* in *Sa. islandicus*; Appendix Figure S2), further distinguishing the representatives of the class 5 OLD enzymes from the four previously defined OLD classes (Akritidou & Thurtle-Schmidt, 2023; Cheng *et al*., 2021; Huo *et al*, 2024; Li *et al*, 2024; Oh *et al*, 2023). To study whether *Sulfolobales* OLD-family proteins are implicated in anti-virus defense, we overexpressed Cran1 alone or in combination with the putative accompanying membrane protein SiRe_0087 in *Sa. islandicus* E233S and challenged the recombinant cells with Sulfolobus tenchongensis spindle shaped virus 2 (STSV2) and Sulfolobus monocaudavirus 1 (SMV1) using a spot assay. Neither of the Cran1 overexpression strains showed detectable antiviral ability. On the contrary, the strain overexpressing Cran1 alone or in combination with the membrane protein SiRe_0087 were more sensitive to STSV2 infection than the control strain (Figure EV3A). Similarly, overexpression of Cran1 alone or together with SiRe_0087 in *Escherichia coli* MG1655 did not provide detectable protection against bacteriophages T1, T2, T3, T4, T5, and T7 under the assay conditions tested (Figure EV3B). Although we cannot exclude the possibility that the antiviral activity of Cran1 is specific towards particular viruses (which were not among those tested in this work), our results suggest that Cran1 does not function in antiviral defense but may have other functions.

### Cell cycle-dependent expression of Cran1 appears to be essential for cell viability

We have previously observed that overexpression of aCcr1, a transcription factor that regulates the cell cycle progression in *Sa. islandicus* REY15A by suppressing the expression of a gene encoding a key cytokinesis protein CdvA, resulted in 4-fold transcriptional repression of *cran1* (Yang *et al*., 2023b). Analysis of the promoter region of *cran1* revealed the presence of the aCcr1 recognition sequence (Yang *et al*., 2023b). Electrophoretic mobility shift assays showed that purified aCcr1 binds to the oligonucleotides encompassing the promoter region of *cran1* (P*_cran1_*; Appendix Figure S3), confirming that aCcr1 regulates the transcriptional expression of Cran1.

To analyze whether Cran1 exhibits cell cycle-dependent expression at the protein level, we constructed a *Sa. islandicus* REY15A strain in which the Cran1 is expressed *in situ* with a Flag-tag at the C-terminus (Appendix Figure S4A). We verified that the growth and cell morphology of the *in situ* Flag-tagged Cran1 strain are not different compared to those of the wild type (Appendix Figure S4BC). Then, the cell cycle of the *in situ* Flag-tagged strain was synchronized with 6 mM acetic acid (Figure 4A), and the expression levels of Cran1 at different time points were examined by Western blotting with anti-Flag antibody. As shown in Figure 4B and C, Cran1 exhibited a cell cycle-dependent expression pattern, with the protein levels being the highest during the M-G1 and S cell cycle phases (i.e., 3-6 h following the removal of acetic acid; Figure 4B). The cyclic expression of *Cran1* at the transcriptional level was further confirmed by RT-qPCR using synchronized *S. islandicus* E233S cells (Figure 4D), fully recuperating our previous transcriptomic result (Yang *et al*., 2023b). Notably, during the logarithmic growth phase, the amount of Cran1 in the cells was about five times higher than during the stationary phase (Figure 4E).

**Figure 4.**
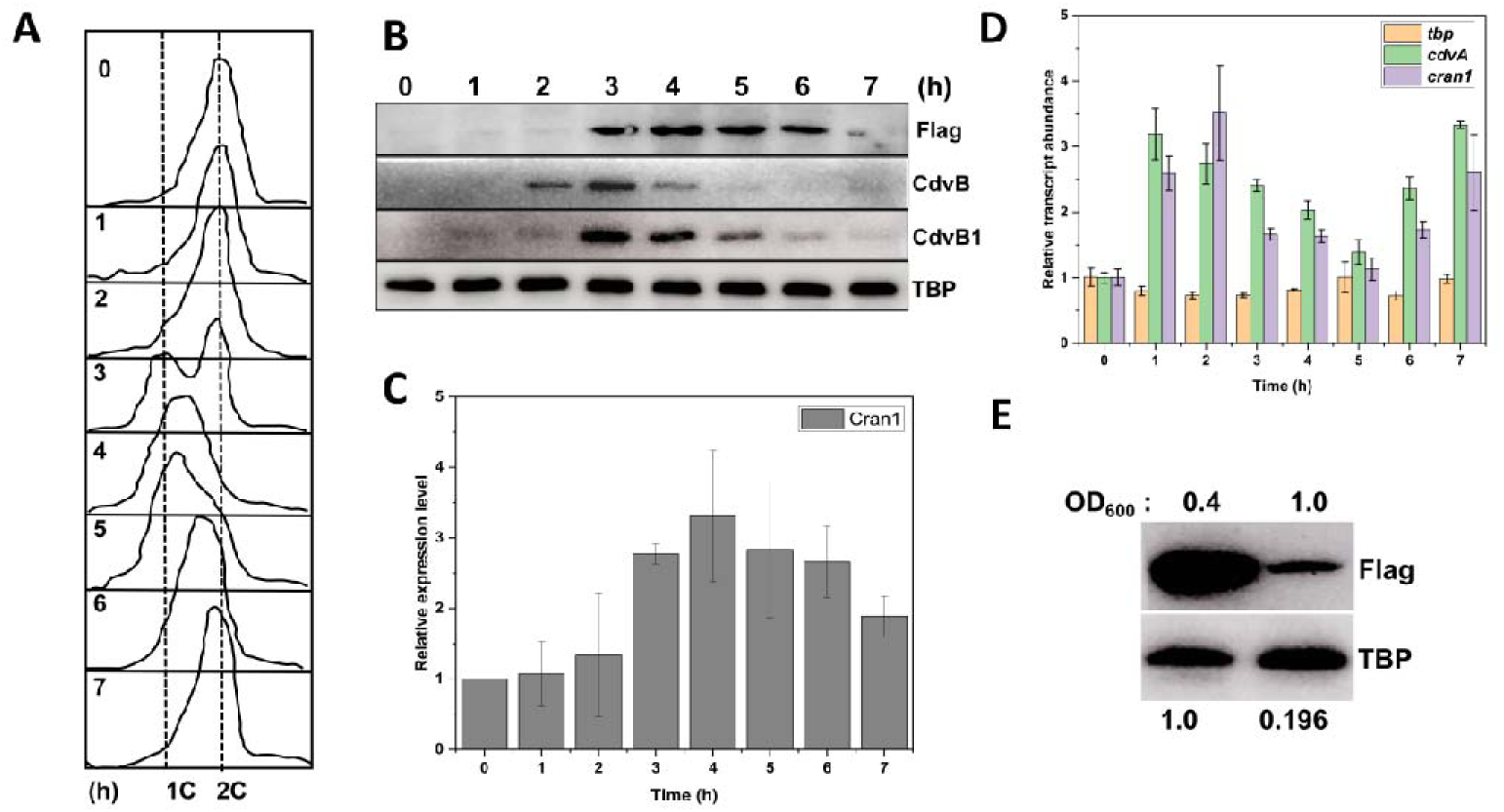
The cyclically expressed gene *cran1* is essential for cell viability. (**A**) Flow cytometry profiles of samples of a synchronized Cran1 in-situ-flag strain in one cell cycle. The cells were synchronized at G2 phase after treated with 6 mM acetic acid for 6 h before released by removing the acetic acid. The cultures were collected at different time points (0, 1.0, 2.0, 3.0, 4.0, 5.0, 6.0 and 7.0 h) and subjected to flow cytometry analysis. Cells started to divide at 2.0h as shown by the appearance of cells with one copy chromosome (1C) and the ratio of dividing cells reached the highest level at about 3-3.5h. As the cell cycle proceeded, cells with two copy chromosomes (2C) became dominant at 7h. (**B**) Changes of the protein levels of Cran1 during a cell cycle detected by Western blotting using anti-Flag antibody. The known periodic expression of CdvB and CdvB1 were used as positive controls, and TBP (TATA-box binding protein) was used as a loading control. (**C**) Quantification the result in (B) (3 biological replicates). (**D**) RT-qPCR detection of transcriptional levels *cdvA* and *cran1* during a cell cycle (3 biological replicates). (**E**) Western blotting to the protein levels of Cran1 during exponential (OD_600_=0.4) and stationary (OD_600_=1.0) phases, respectively.

To assess the importance of Cran1 for the cell, we attempted to knock out *Cran1* using the endogenous CRISPR-based genome editing method in *Sa. islandicus* REY15A (Li *et al*., 2016; Wei & Li, 2023). However, all attempts (at least five times) failed to yield a viable knockout clone, suggesting that *Cran1* is an essential gene. The apparent essentiality of Cran1 contrasts the dispensability of defense systems, including those that have OLD family components in bacteria, and suggests that Cran1 plays an important role for the survival of the cell even in the absence of virus infection. Notably, unlike *cran1*, the gene (*sire_0087*) which forms a putative operon with *cran1* (Appendix Figure S2) and encodes a hypothetical membrane protein could be knocked out, and the knockout strains exhibited growth and morphology phenotypes similar to those of the wild type (Appendix Figure S5). However, our attempts to knock out the whole operon was unsuccessful, further suggesting that the role of Cran1 is different from that of bacterial OLD proteins.

To further assess the importance of Cran1 for the cell, we successfully constructed a *Cran1* knockdown strain using the endogenous CRISPR-Cas based method (Peng *et al*, 2015; Zink *et al*, 2019). As shown in Figure EV4AB, the knockdown strain with four spacers targeting *cran1* had reduced protein level (around 50%) and exhibited apparent growth retardation. Microscopy analysis revealed that the Cran1 knockdown strain generated more (5.6%, n=504) large cells (approximately 2 µm in diameter) when compared to the control strain (0.55%, n=720) (Figure EV4C). Consistent with the growth retardance phenotype, flow cytometry analysis showed that there was slight difference, although not significant, between the knockdown and the control strains, with smaller population of 1C cells in the knockdown strain than in the wild type strain (Figure EV4D), suggesting a slowdown of DNA synthesis or impairment of chromosome segregation and/or cell division. The relatively mild phenotype obtained with the knockdown strain may be linked to the limited efficiency of the transcript depletion, i.e., cells with 50% of the native Cran1 may function relatively normally. Nevertheless, these results further suggest that Cran1 plays an important role in *Sa. islandicus* cell cycle.

### Overexpression of Cran1 variants with the wild type ATPase domain impairs the cell cycle progression

Strikingly, overexpression of Cran1 resulted in dramatic growth retardation (Figure 5A), which was associated with a considerable increase in cell size, most noticeably after 24 hours post-induction (average diameter of 4.54 µm; n=100) compared to the control cells (∼1.2 µm), with 95% of the cells exhibiting 50-fold increase in volume (Figure 5B and 5C). Additionally, flow cytometry analysis showed an increase in the number of genome equivalents within the population of large cells (Figure 5D). All these phenotypes imply that Cran1 is involved in cell cycle progression. By contrast, overexpression of SiRe_0087 had no effect on the growth and cell morphology (Appendix Figure S5BCD). Co-expression of Cran1 and SiRe_0087 yielded phenotypes generally similar to expression of Cran1 alone (Appendix Figure S5BCD). These results suggest that although *sire_0087* and *cran1* are putatively located in the same operon, their functions may not be directly linked or correlated. To analyze whether the cell cycle-related function of Cran1 is conserved in other members of the order Sulfolobales, we overexpressed a Cran1 ortholog from *S. acidocaldarius* DSM639 (Sac Cran1) in *S. islandicus* E233S. Sac Cran1 overexpression also resulted in growth retardation, larger cell size, and increased cellular DNA content (Figure EV5), suggesting that the cell cycle-related function of Cran1 is conserved in Sulfolobales, consistent with the conservation of gene neighborhood of *cran1* in other members of the order (Figures 1B and Appendix Figure S2).

**Figure 5.**
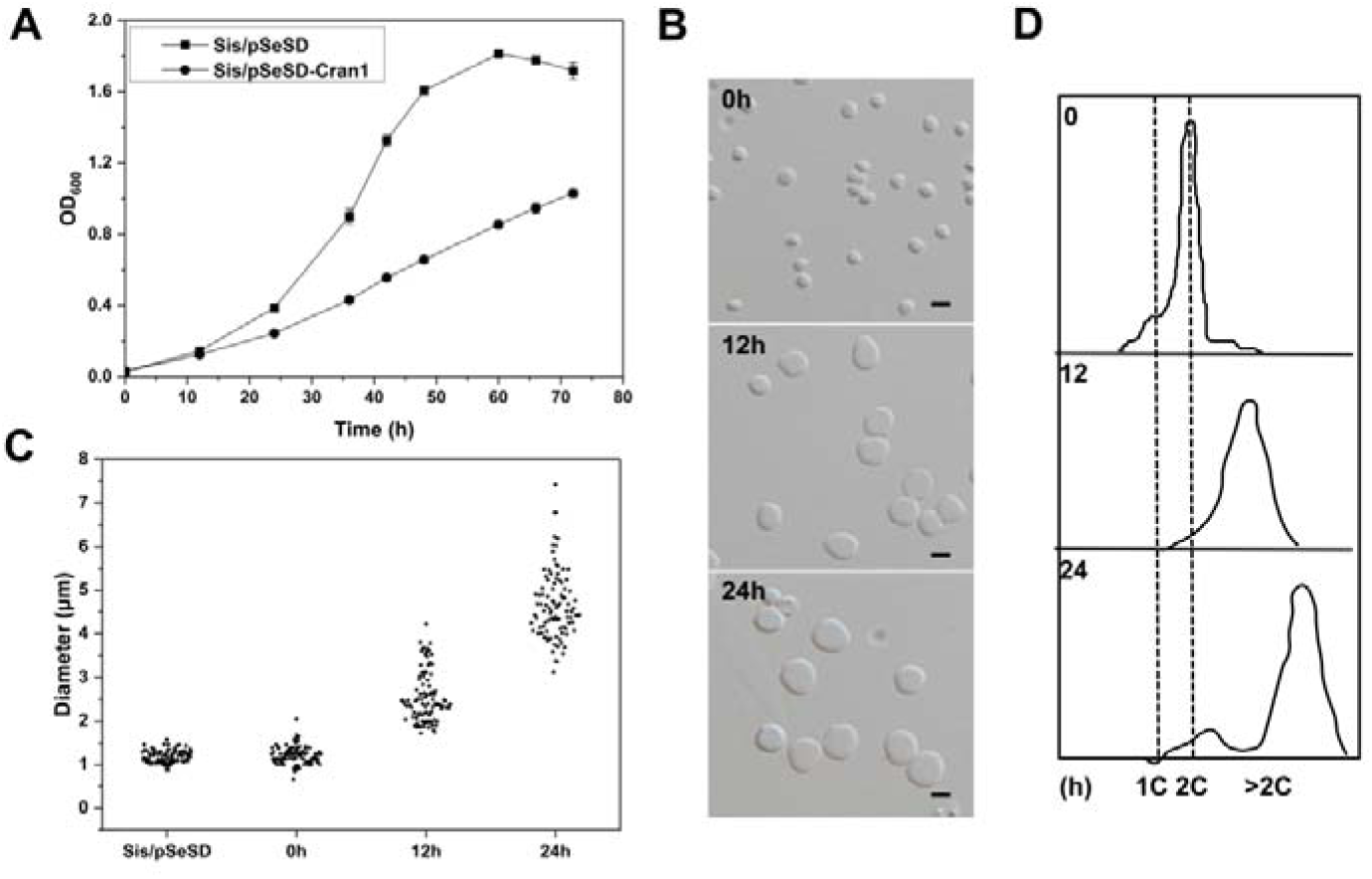
Overexpression of Cran1 leads to retarded cell growth and cell enlargement. (**A**) Growth curves of cells over-expressing the C-terminal His-tagged Cran1. The cells were inoculated into 30 ml induction medium ATV to a final estimated OD_600_ of 0.03. The growth was monitored at OD_600_. Each value was based on data from three independent measurements. Cells harboring the empty plasmid pSeSD were used as a control. (**B**) Bright-field microscopy (DIC) of cells over-expressing Cran1. Cells cultured in the induction medium were taken at different time points and observed under a NIKON-E microscope. Upper, middle, and lower panels, 0, 12, and 24h. Scale bars: 2 μm. (**C**) Cell size statistics of the data in (B). Cell cultures were sampled at the indicated time points and observed under the microscope. The diameters of ∼100 cells were measured using ImageJ software for each culture repeat. (**D**) Flow cytometry analysis of the DNA content of Sis/pSeSD and Sis/pSeSD-Cran1 cells cultured in MATV medium.

In order to explore whether the enzymatic activities of Cran1 are important for the observed effects, we constructed *Sa. islandicus* strains overexpressing different ATPase active site mutants and tested their phenotypes. Overall, the effects of different mutants on the culture growth rates and cell morphology were consistent with the *in vitro* ATPase activity results. In particular, the strains expressing the ATPase mutants that had the strongest impacts on the ATP hydrolysis *in vitro* approached the phenotypes of control cells containing the empty vector, whereas mutants in which the ATPase activity was only mildly affected, resembled the strain overexpressing the wild type Cran1 and displayed growth retardation and increase in cell size. In the strain overexpressing Cran1(T37A), a mutant with severely compromised ATPase activity *in vitro*, the majority of cells (but not all) were of normal size, and their growth rate was closest to that of the strain carrying the empty vector pSeSD (Figure 6A). The strains expressing Cran1(H280A) and Cran1(S281A) also had improved growth rate compared to the cells overexpressing the wild type Cran1 (Figure 6A). Notably, the Cran1(S281A) cells were enlarged and more irregular, with bud-like formations and dumbbell-like morphologies (Figure 6E), compared to the other mutant cells that were uniformly enlarged and spherical. Similar abnormal cell morphologies were observed in the case of the cell division impaired *Sa. islandicus* strain overexpressing the dominant-negative mutant of the key cytokinetic ATPase, Vps4 (Liu *et al*, 2025b). It is noteworthy that in the class 1 and 2 OLD proteins, mutation of the key aspartic acid residue in the Walker B motif results in a complete loss of the ATPase activity. In Cran1, the Walker B motif residues E246 and E247 are predicted to coordinate a magnesium ion. However, in our experiments, substitution of either of the two residues with alanine had no effect either in *in vitro* or *in vivo*. Analysis of the structural model suggested that T37 is closer to the magnesium ion than E246 and E247, and may play a more important role in magnesium ion coordination or in maintaining an functional conformation compared to E246 and E247, which may explains why mutation of either Walker B residue had no significant effect on the ATPase activity (Figure 3D) and still led to growth retardence (Figure 6B).

**Figure 6.**
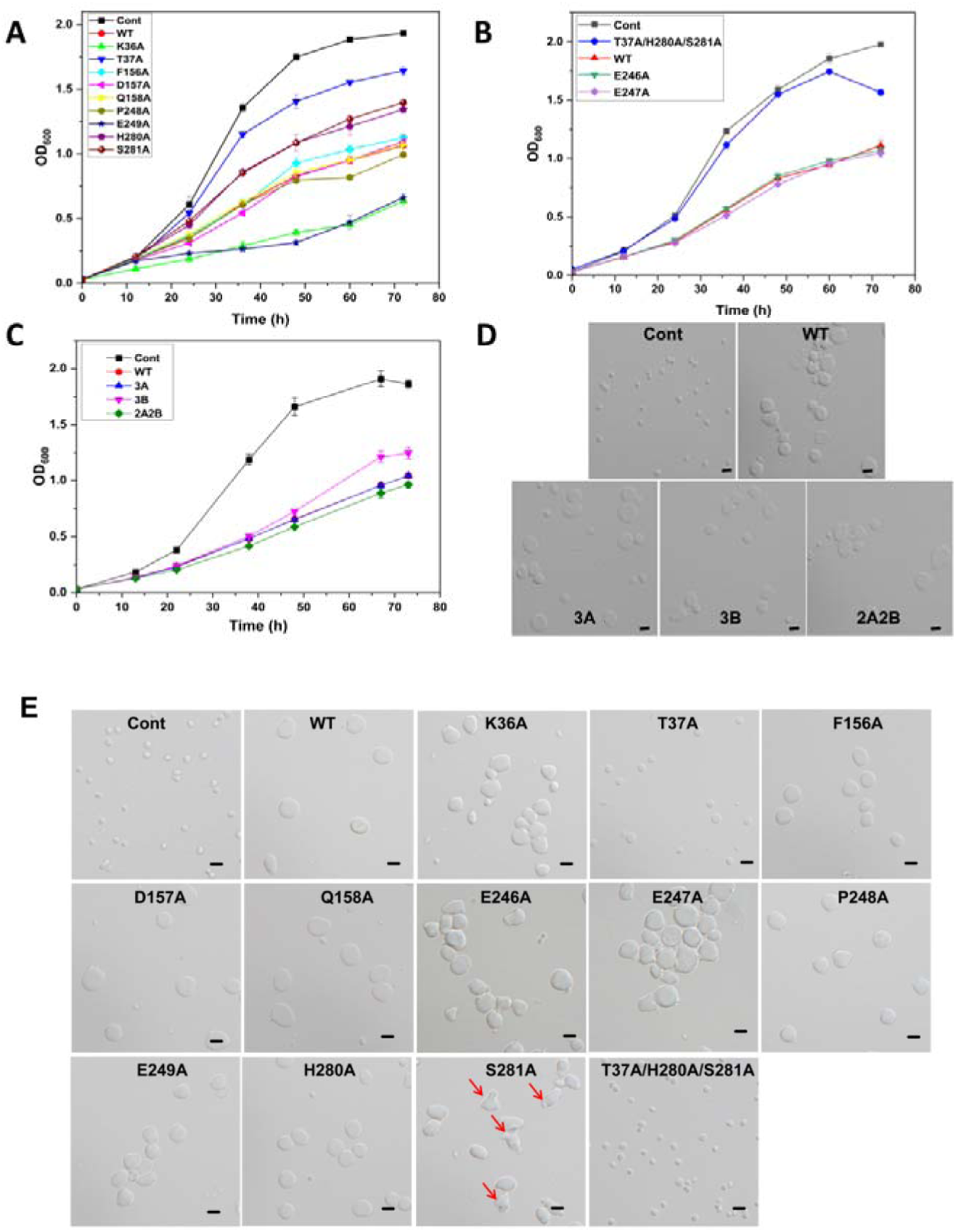
The ATPase activity of Cran1 is vital for cell cycle progression. (**A**) Growth curves of the single predicted key amino acid residue mutant overexpression strains for the ATPase activity. (**B**) Growth curves of additional predicted key amino acid residue mutants E246A and E247A, and a multiple mutant T37/H280/S281 for the ATPase activity. (**C**) Growth curves of the putative metal ion binding site mutant overexpression strains for the nickase activity. “3A” denotes E338A/L391A/D393A; “3B”, E341A/E409A/E411A; “2A2B”, the L391A/D393A/E409A/E411A. (**D**) and (**E**) Representative images of cells of the nickase deficient mutant overexpression strains in (C) and the predicted key amino acid residue mutant overexpression strains for the ATPase activity in (A-B), respectively. Samples were taken from arabinose induction for 24h. Scale bars, 2 μm.

Given that none of the single mutants completely recovered the phenotype of control cells, we constructed a strain overexpressing Cran1(T37A/H280/S281A), a triple mutant, which lost both the ATPase and nuclease activities *in vitro* (Figure 3D). Overexpression of this mutant *in vivo* yielded cells with morphology identical to that of the control cells (Figure 6B and 6E). Unexpectedly, *Sa. islandicus* strains carrying plasmids overexpressing any of the nuclease active site mutants, which abrogate the nuclease activity *in vitro*, showed the same phenotypes as those observed in the strain overexpressing the wild-type Cran1 (Figure 6C and 6D). We verified that the protein expression in all mutants was at similar levels using Western blotting, confirming that the phenotypic difference was not caused by differences in protein expression level (Figure EV2F). Collectively, our results suggest that an inteact ATPase domain is required for the Cran1 variants to interfere with the normal progression of the cell cycle.

### Overexpression of Cran1 blocks cytokinesis

To analyze which of the cell cycle phases was impacted by the overexpression of Cran1, we synchronized the Cran1 and Cran1(T37A/H280A/S281A) overexpression strains to G2 phase, with cells harboring the empty vector pSeSD used as a control (Figure 7). To ensure sufficient protein concentration, arabinose was added to induce protein expression two hours before acetic acid removal. The control cells started to divide ∼4 h following the release of the cell cycle arrest. Profiles similar to those of the control cells were obtained for the strain overexpressing the inactive Cran1 ATPase mutant Cran1(T37A/H280A/S281A) (Figure 7B). By contrast, the division of the wild type Cran1 overexpression strain was blocked. Notably, although cell cycle could not progress into the cell division phase, genome replication continued in the cells expressing the wild type Cran1 leading to accumulation of intracellular DNA, suggesting that the cell cycle was blocked in the S phase.

**Figure 7.**
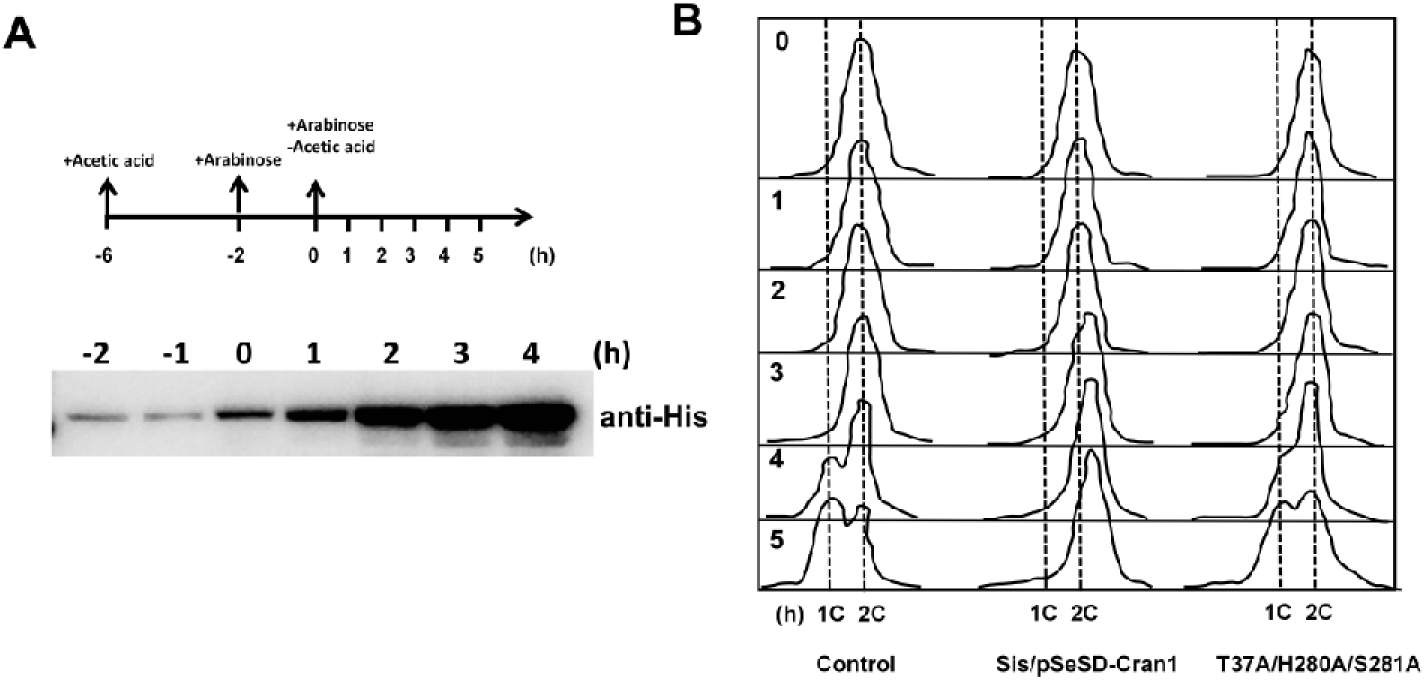
Overexpression of Cran1 blocks the process of cell division. (**A**) A schematic showing cell synchronization and induction of Cran1 over-expression with arabinose (0.2%). The time for the acetic acid treatment and arabinose induction are indicated. E233S containing the empty plasmid (Sis/pSeSD) was used as a control. (**B**) Flow cytometry profiles of DNA content distribution of cells 0-5 hrs after acetic acid removal.

### Cran1 may be involved in chromatin processing related activities in M-G1 and S phases

To gain further insights into how Cran1 functions, we determined the subcellular localization of Cran1 by chromatin fractionation. As shown in Figure 8A, a fraction of Cran1 was associated with the chromatin in the logarithmic growth phase, whereas almost all of the protein was located in the cytoplasm during the stationary phase (Figure 8A). To assess whether Cran1 is involved in chromatin associated activities, we performed pulldown assay using genome coded Flag-tagged Cran1 as a bait, followed by identification of the purified proteins by mass spectrometry. Interestingly, multiple chromatin-associated proteins co-purified with Cran1, with Cren7 and histone deacetylase superfamily protein (SiRe_1988) being the most abundant (Figures 8B and 8C, Appendix Table S4). Notably, both Cran1 and Cren7 are among the essential genes regulated by aCcr1 (Yang *et al*., 2023b). These results suggest that Cran1 likely functions in cooperation with chromatin proteins.

**Figure 8.**
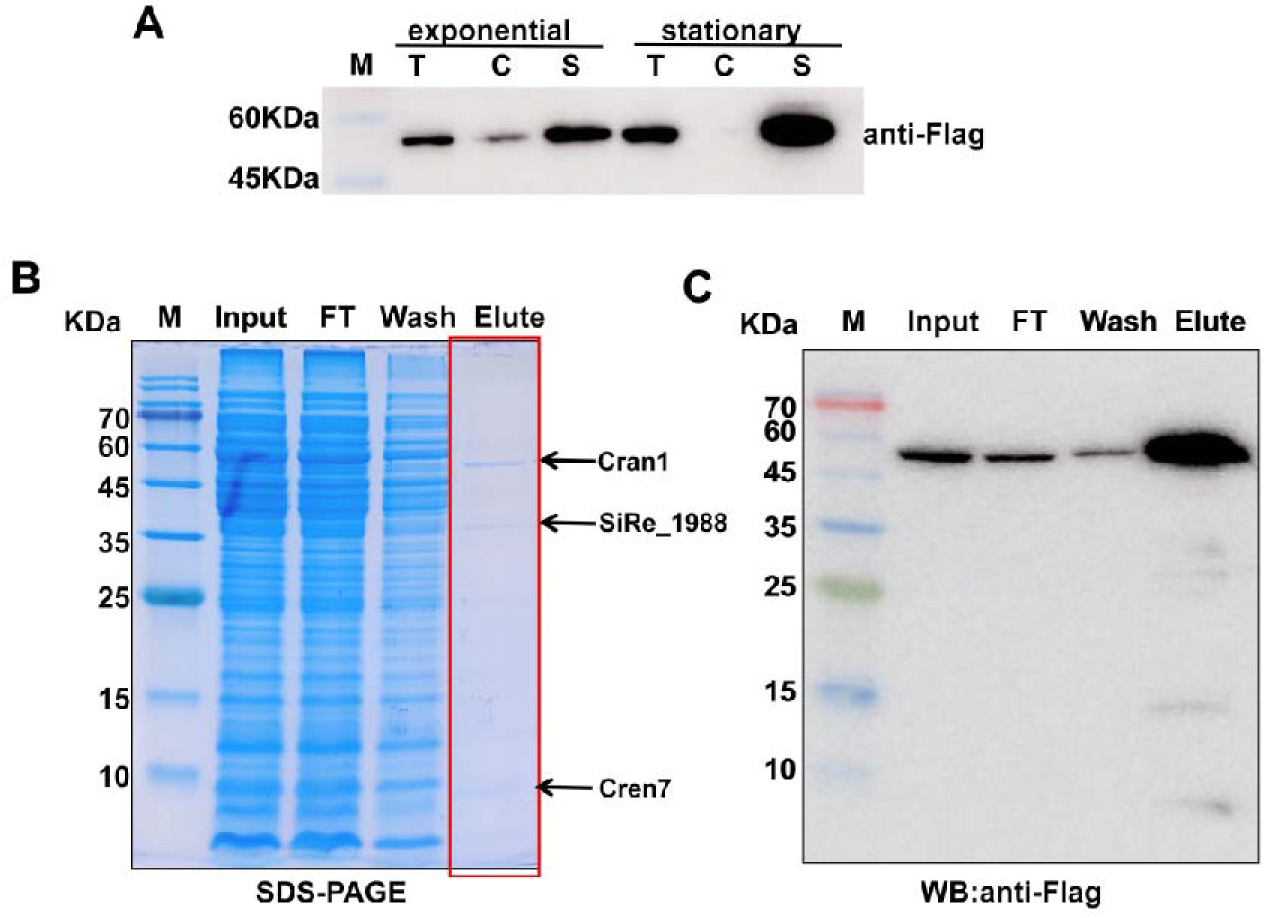
Subcellular localization of Cran1. (**A**) Cellular localization of Cran1 in the cytoplasm or chromatin of cells during the exponential phase (OD_600_=0.4) of cell growth as well as in stationary cells (OD_600_=1.0). T, total protein. C, Chromatin fractions. S, Soluble fractions. (**B**) SDS-PAGE and (**C**) Western blotting of immunoprecipitation of Cran1 *in-situ* Flag strains. Input represents cell lysate, FT indicates flow-through, Wash indicates washing of magnetic beads using buffer A to remove nonspecifically bound fractions and Elute indicates elution fractions of target proteins.

## DISCUSSION

Here, we characterized a representative of a new class of OLD family proteins, class 5, and the first OLD family protein from archaea. Contrary to our initial expectation, Cran1 did not appear to participate in the anti-virus defense but was rather involved in the cell cycle progression. Consistently, unlike in the case of bacterial defense-related OLD proteins, which are not essential for the normal cell growth or viability (Akritidou & Thurtle-Schmidt, 2023), the *cran1* gene of *Sa. islandicus* could not be deleted. Cran1 displays cyclic expression, with the protein level being high during the progression of the cell cycle from the M-G1 into the DNA synthesis (S) phase, and being depleted during the post-replicative G2 phase (Figure 4B). Importantly, Cran1 is enriched in the chromatin fraction in vigorously growing cells and immunoprecipitation showed that Cran1 is associated with the chromatin protein Cren7 and a histone deacetylase homolog. Overexpression of Cran1 does not affect genome replication but appears to impair chromosome segregation and/or cell division. Collectively, these observations suggest that Cran1 functions during the M-G1 and S phases and is involved in the chromatin processing related activities, although the possibility that it plays a role in antiviral defense under particular circumstances or against specific viruses can not be formally excluded.

The knockdown of Cran1 gene resulted in only 50% depletion of the corresponding protein. Although the strain displayed mild but notable changes in the growth and cell division phenotypes, the corresponding cells were still able to complete the cell cycle progression. By contrast, the cell cycle was strongly impaired in the Cran1 overexpression strain, with the cells displaying an increase in size and chromosomal copy numbers, suggesting blockage of cytokinesis. We hypothesize that during the normal cell cycle progression, Cran1 levels need to be reduced during the G2 phase prior to cell division (Figure 4B), when chromosomes undergo condensation in preparation for segregation. This is apparently achieved through a combination of transcriptional regulation by aCcr1 and degradation of the protein (Risa *et al*., 2020; Yang *et al*, 2023a). It has been reported that the half-life of the *SSO2277* mRNA, a close ortholog of Cran1 from *Saccharolobus solfataricus*, is only 3 min (Andersson *et al*, 2006). Furthermore, there was a slight accumulation of SacCran1 (Cran1 from *Sulfolobus acidocaldarious*) when the cells were treated with the proteasome inhibitor Bortezomib (Risa *et al*., 2020). These observations imply that the mRNA of Cran1 is tightly controlled at the transcriptional level and the level of Cran1 could be further regulated at the protein level by proteasome degradation. The dual regulation that Cran1 is subjected to *in vivo* emphasizes the importance of cell cycle phase-dependent functioning of this protein.

The exact role of Cran1 during the cell cycle progression remains unclear. However, given that Cran1 expression is elevated following cell division, during the M-G1 and S phases, we hypothesize that Cran1 facilitates genome replication by relaxing the chromosome using its nickase activity. By contrast, overexpression of Cran1 beyond the S phase would be expected to interfere with the chromosome condensation and subsequent cytokinesis. Notably, the inability to undergo cell division does not appear to prevent the subsequent round of genome replication, further suggesting that genome replication and cytokinesis are not strictly coupled in *Sa. islandicus*, consistent with previous results (Liu *et al*., 2025b). This could explain the emergence of large cells with multiple copies of the genome. This result is consistent with the more efficient proliferation of the STSV2 virus in the Cran1 overexpression strain. Indeed, similar to some other viruses, STSV2 is known to block the cell division, arresting the cell cycle in the DNA synthesis phase, which ensures the access to the host’s replication machinery (Fan *et al*, 2018; Kiro *et al*, 2013; Liu *et al*., 2021a; Stewart *et al*, 2013). Cran1 protein level during the exponential growth phase is five times higher than during the stationary phase, consistent with the possibility that Cran1 promotes genome replication during the rapid growth phase.

In eukaryotes, histone modification is an important mechanism for regulating genome accessibility (Bannister & Kouzarides, 2011). Immunoprecipitation experiments showed that Cran1 is associated with histone deacetylase homolog and Cren7, Sac7d/Sso7d, SSB (single-stranded DNA binding protein), and Alba (Appendix Table S4). Chromatin proteins have various post-translational modifications (acetylation, methylation, phosphorylation, etc.)(Wolffe & Hayes, 1999), which affect not only the properties of the proteins themselves, but also the structure of the chromatin and gene expression. For example, many chromatin proteins of *Sa. islandicus* have been reported to be post-translationally acetylated or methylated (Cao *et al*, 2019; Vorontsov *et al*, 2016). Therefore, chromatin proteins and their post-translational modifications are likely to play an important role in the cell cycle progression. Given that Cran1 has active ATPase and nickase activities, we hypothesize that it plays a role in chromatin relaxation and/or unwinding of supercoiled DNA in cooperation with chromatin proteins, such as Cren7, during the S phase, facilitating the replication and subsequent chromosome segregation. However, the mechanism underlying the putative involvement of Cran1 in chromatin processing remains unknown.

Structural and biochemical studies have greatly increased our understanding on the anti-phage functions of the OLD family proteins in bacteria in recent years. In the Gabija antiviral system, GajA, a representative of the Class 2 OLD family, forms a tetramer, with the ATPase domains clustered at the center and the Toprim domains located peripherally (Li *et al*., 2024). ATP binding in the ATPase domain stabilizes the insertion region within the ATPase domain, keeping the Toprim domain in a closed state. Upon ATP depletion by phages, the Toprim domain opens to bind and cleave the DNA substrates (Li *et al*., 2024).Notably, GajA has likely lost its ATP hydrolysis activity during evolution but retained the nucleotide-binding capacity (Cheng *et al*., 2021), with the ATPase domain playing a regulatory role, stimulating the nuclease activity in response to viral invasion (Cheng *et al*., 2023; Cheng *et al*., 2021). Similarly, the ATPase domains of the Class 2 OLD proteins BpOLD (*Burkholderia pseudomallei* OLD) and XccOLD (*Xanthamonas campestris pv. Campestris* OLD), which catalyze the nicking and cleavage of DNA substrates, are thought primarily to promote oligomerization and control how and when the nuclease domain gains access to the substrate (Schiltz *et al*., 2019). We found that deletion or inactivation of the Cran1 ATPase domain abolishes the nuclease activity, suggesting that there is a cross talk between the two domains, similar to what has been reported to other OLD family proteins. Given that the presence or absence of ATP, dATP, ADP, and AMP did not affect the nuclease activity of Cran1 *in vitro*, we presume that the proteins were copurified with the bound nucleotide. It is tempting to speculate that it is the inherent crosstalk between the sensor ATPase domain and the effector nuclease domain that predisposed the OLD family enzyme for recruitment as an important player in the cell cycle control in Sulfolobales. Indeed, as evident from the *in vivo* experiments with the strain overexpressing the nucleotide binding and ATPase deficient mutant (T37A/H280A/S281A) which did not show growth inhibition, the ATPase domain of Cran1 is crucial for its function *in vivo*. In conclusion, our study has revealed an unexpected novel function of an archaeal OLD family protein in cell cycle progression.

## Supporting information

Appendix Figures and Tables.docx

## DATA AVAILABILITY

All data supporting the findings of this study are available within the article and its Supplementary Information, or from the corresponding author upon reasonable request.

## Acknowledgements

We would like to thank members of the CRISPR and Archaea Biology Research Center for helpful discussions and technicians from the Core Facilities for Life and Environmental Sciences, State Key Laboratory of Microbial Technology of Shandong University for assistance. This work was supported by National Natural Science Foundation of China [32393973 and 32370033, the State Key Laboratory of Microbial Technology Open Projects Fund (Project NO. M2023-20) to Y.S. and Postdoctoral fellowship Program of CPSF (GZC20231471) and Postdoc Innovation Project of Shandong Province (SDCX-ZG-202400122) to Y.Y. M.K. was supported by a grant from the Agence nationale de la recherche (ANR-23-CE13-022).

## Author contributions

**Yunfeng Yang**: Conceptualization; Data curation; Formal analysis; Investigation; Visualization; Methodology; Writing-original draft and editing. **Shikuan Liang**: Investigation; Methodology. **Junfeng Liu**: Conceptualization; Visualization; Methodology. **Xiaofei Fu**: Investigation. **Pengju Wu**: Methodology. **Haodun Li**: Methodology, **Jinfeng Ni**: Writing-review and editing. **Qunxin She**: Conceptualization; Writing-review and editing. **Mart Krupovic**: Conceptualization; Phylogenetic analysis; Writing-review and editing. **Yulong Shen**: Conceptualization; Supervision; Funding acquisition; Project administration; Writing-review and editing.

## Expanded View Figures

**Figure EV1.**
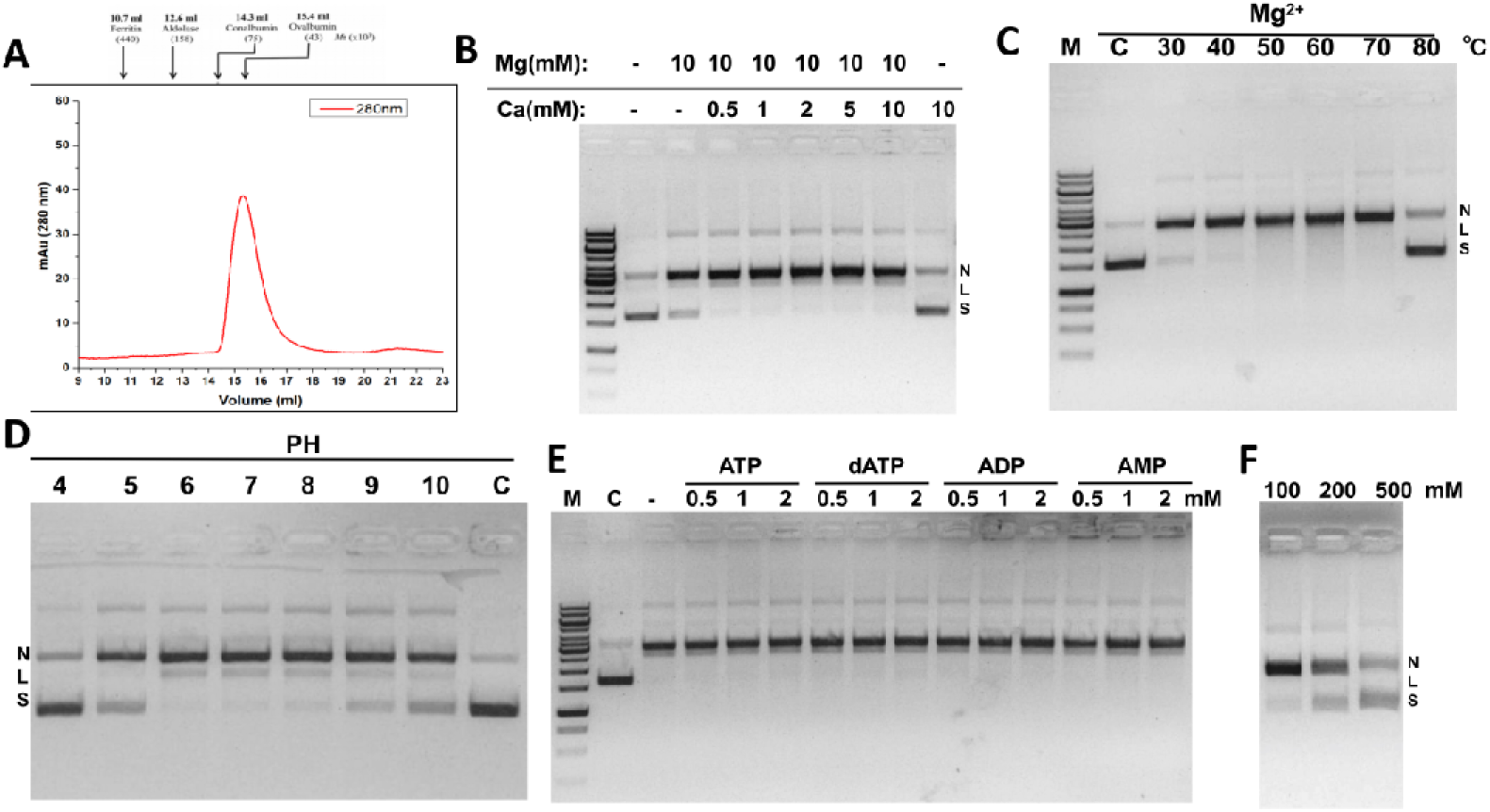
Biochemical properties of Cran1 in vitro. (**A**) Size exclusion profile of the purified wild-type Cran1. The protein was expressed in E. coli BL21 and purified by heat treatment, nickel affinity, and gel filtration with a Superdex 200 column as described in the “Materials and Methods”. Cran1 was assayed for nickase activity under different temperature gradient (**B**) as well as PH gradient conditions (**C**). C refers to the control reaction without Cran1 at pH 7 and 37LJ.In these reactions, 300 ng of PUC19 DNA was incubated with 2 µM Cran1 in a final volume of 20µl. Reactions were performed at 70°C for 30 min and then stopped by the addition of 4 µl of 6×loading dye containing 20 mM EDTA. Samples were analyzed via native agarose gel electrophoresis. (**D**) Gradual increase in the concentration of calcium ions under 10 mM magnesium ions to observe the effect of two-metal ion-catalyzed conditions on the activity of Cran1 nickase. (**E**) Observation of the effect of 0.5, 1, and 2 mM of ATP, dATP, AMP, and ADP on the activity of Cran1 nickase. (**F**) Effect of KCl at 100, 200, and 500 mM on Cran1 activity. ‘N’, ‘L’, and ‘S’ denote the positions of ‘nicked’, ‘linearized’, and ‘supercoiled’ DNA respectively.

**Figure EV2.**
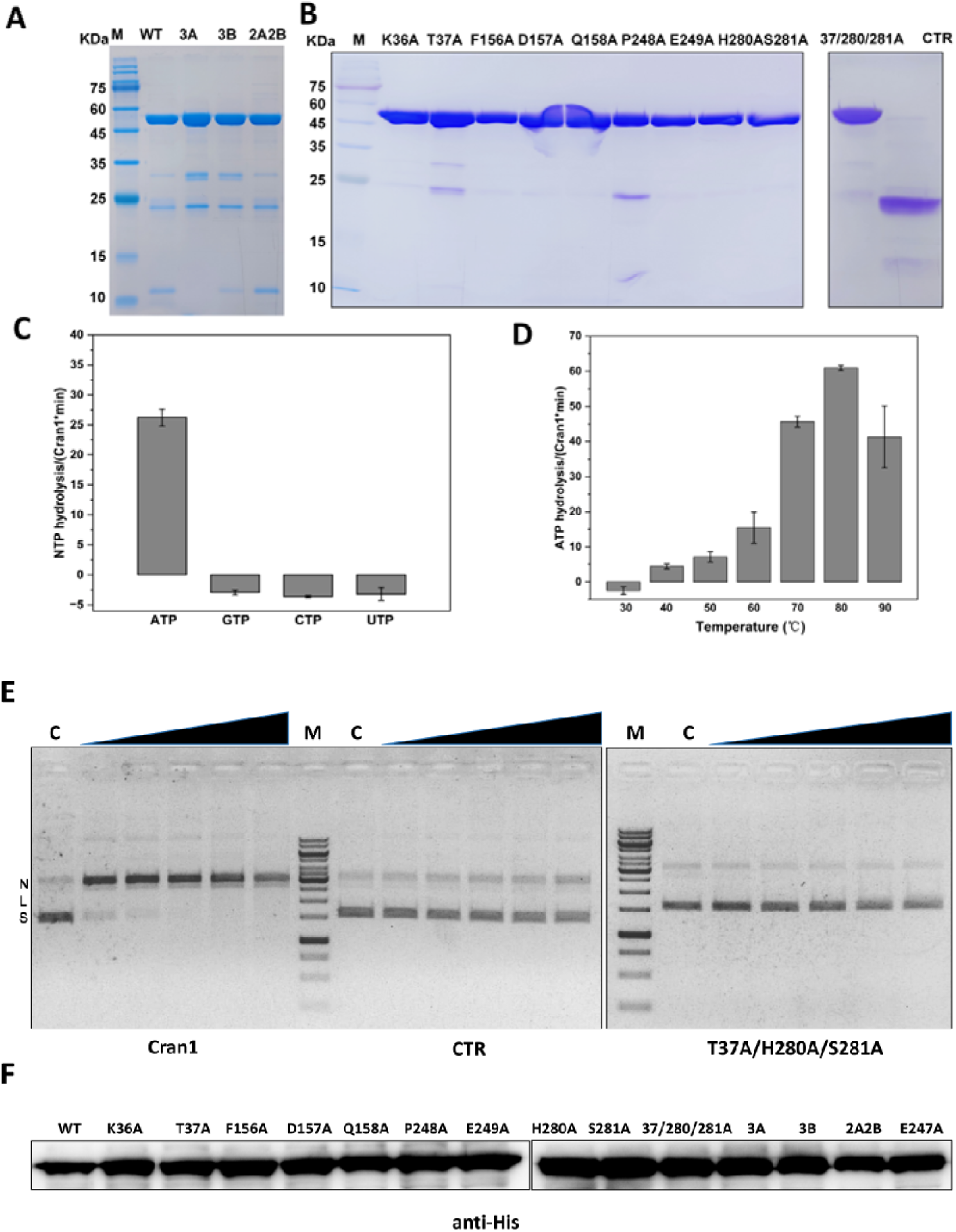
Purification and the activities of the Cran1 wild type as well as mutant proteins. (**A**) and (**B**) SDS-PAGE analysis of the wild type, nuclease active site mutants (A) and the ATPase active site mutants (B) of Cran1. (**C**) Assy of the ability to hydrolyze ATP, GTP, CTP, or UTP. (**D**) The hydrolytic activity of Cran1 on ATP under different temperature conditions. (**E**) The ATPase activity of Cran1 is essential for the performance of nuclease functions. The cleavage experiments of wild-type Cran1 as well as mutant CTR, T37A/H280A/S281A were carried out under the same conditions as in the previous reaction, with magnesium ion conditions at 70°C for 30 min, and protein concentration gradients of 0.25, 0.5, 1, 2, and 4 uM.‘N’, ‘L’, and ‘S’ denote the positions of ‘nicked’, ‘linearized’, and ‘supercoiled’ DNA respectively. (**F)** Western blotting to detect the expression levels of the wild type Cran1 and its mutant overexpression strains. The anti-His tag antibody is used.

**Figure EV3.**
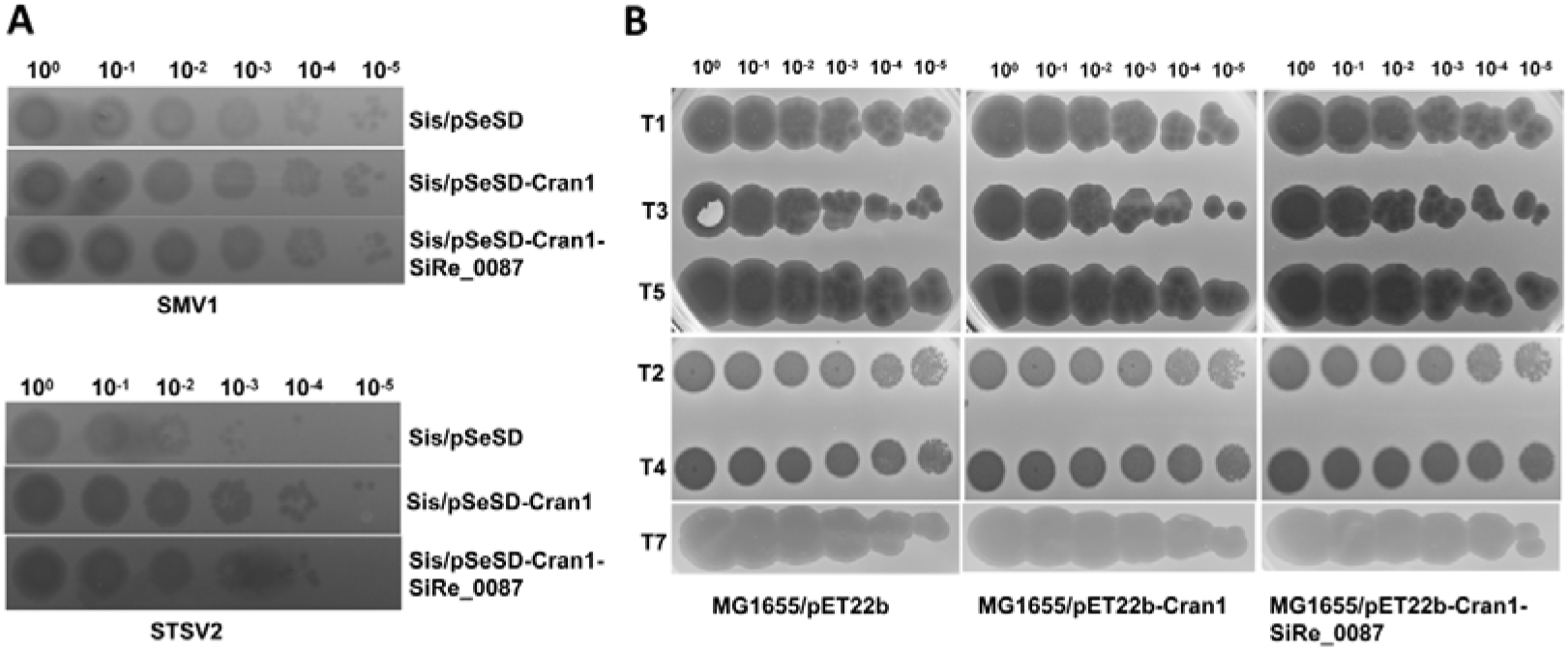
Cran1 is not involved in virus defense. (**A**) Spot assay with the Sa. islandicus virus SMV1 and STSV2 infecting the cells overexpressing Cran1 and cells co-expressing Cran1 and SiRe_0087. (**B**) Spot assay of the infection of phage T1,T2,T3,T4,T5,T7 on Escherichia coli MG1655 cells expressing Cran1 or co-expressing Cran1 and SiRe_0087. Both archaea virus and bacterial phages were diluted in a ten-fold gradient. Strains containing the empty vectors pSeSD and pET22b were used as controls.

**Figure EV4.**
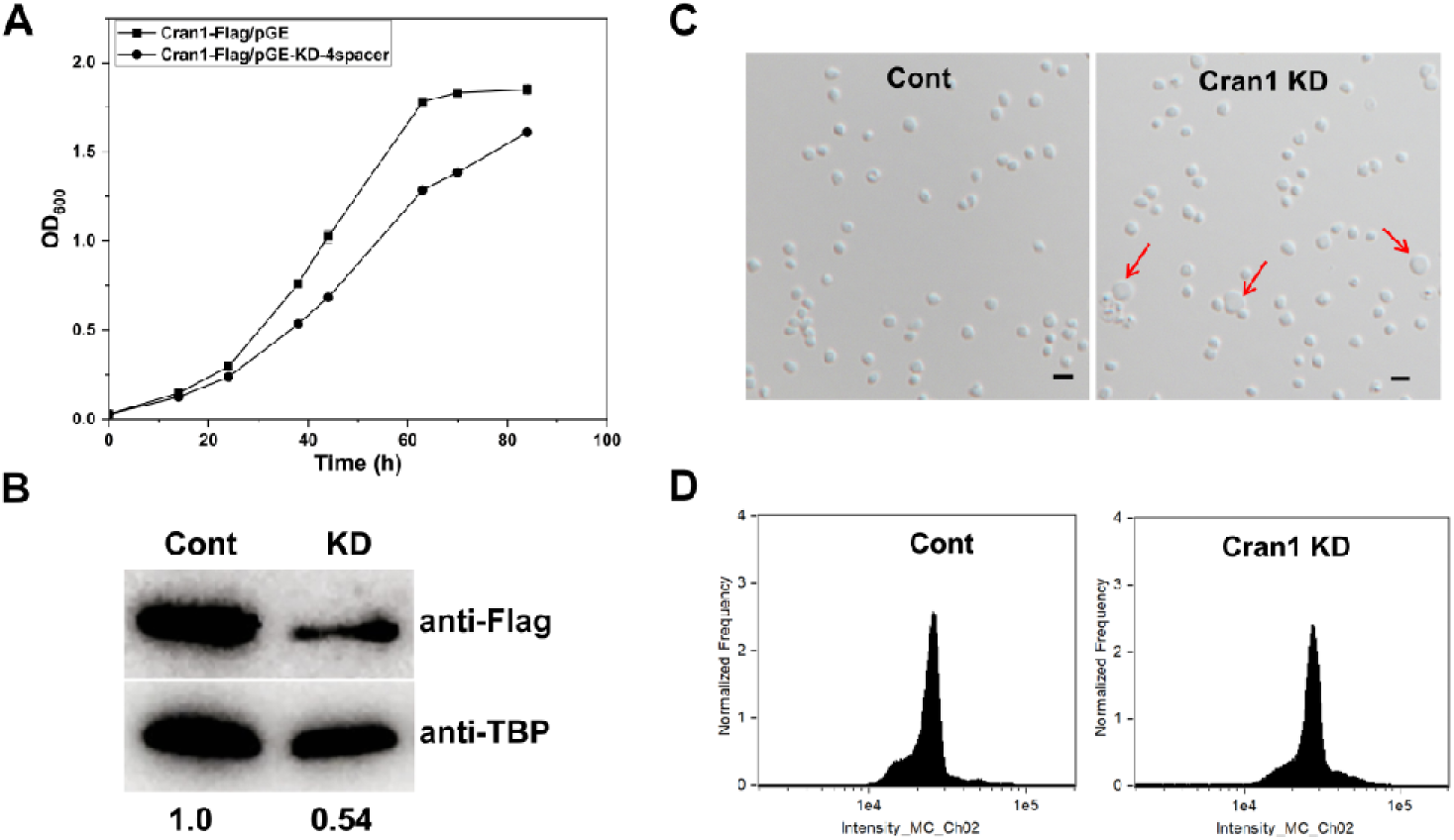
Phenotypic analysis of Cran1 knockdown strains. (**A**) Based on the knockdown technology of endogenous type III CRISPR in Sa. islandicus, an interfering plasmid possessing four tandem repeat spacers was introduced into the Cran1 in situ plus Flag-tagged strains to obtain Cran1 knockdown strains, and the growth curves were determined. (**B**) Western blotting was used to detect protein levels in Cran1 knockdown strains, quantified in gray values using ImageJ software. (**C**) Differential interference contrast (DIC) mode microscopy and (**D**) flow cytometry of Cran1 knockdown strains. Cells cultured in the induction medium ATV were taken at 24h and observed under an inverted fluorescence microscope. DNA content of cells was analysed using an ImageStreamX MarkII Quantitative imaging analysis flow cytometry (Merck Millipore, Germany). Scale bars: 2 μm.

**Figure EV5.**
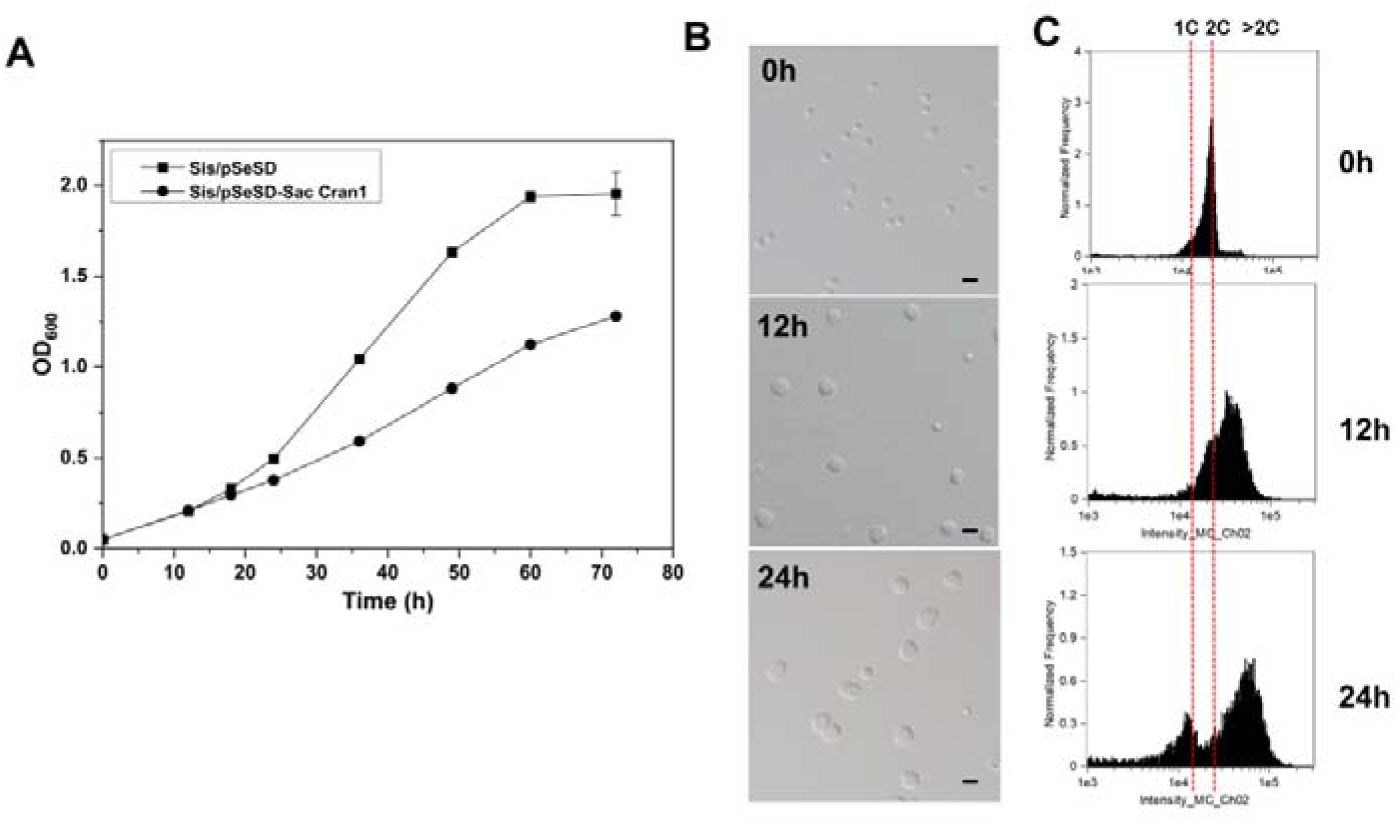
Cells with over-expression of the Cran1 homolog from S. acidocaldarius (Saci_0157, Sac Cran1) has similar phenotypes as those of Cran1. (**A**) Growth curves of strain Sis/pSeSD-Saci_0157. The cells were inoculated into 30 ml ATV medium to a final estimated OD_600_ of 0.03 and the growth was monitored using spectrometer. Each value was based on data from three independent repeats. Cell harboring the empty plasmid pSeSD was used as a control. (**B**) Differential interference contrast (DIC) mode microscopy and (**C**) flow cytometry of cells over-expressing Sac **Cran1**. Cells cultured in the induction medium ATV were taken at different time and observed under an inverted fluorescence microscope. DNA content of cells was analysed using an ImageStreamX MarkII Quantitative imaging analysis flow cytometry (Merck Millipore, Germany). Scale bars: 2 μm.

## References

Akritidou K, Thurtle-Schmidt BH (2023) OLD family nuclease function across diverse anti-phage defense systems. Front Microbiol 14

Altschul SF, Madden TL, Schaffer AA, Zhang JH, Zhang Z, Miller W, Lipman DJ (1997) Gapped BLAST and PSI-BLAST: a new generation of protein database search programs. Nucleic Acids Res 25: 3389–3402

Andersson AF, Lundgren M, Eriksson S, Rosenlund M, Bernander R, Nilsson P (2006) Global analysis of mRNA stability in the archaeon. Genome Biol 7

Antine SP, Johnson AG, Mooney SE, Leavitt A, Mayer ML, Yirmiya E, Amitai G, Sorek R, Kranzusch PJ (2024) Structural basis of Gabija anti-phage defence and viral immune evasion. Nature 625

Aravind L, Leipe DD, Koonin EV (1998) Toprim - a conserved catalytic domain in type IA and II topoisomerases, DnaG-type primases, OLD family nucleases and RecR proteins. Nucleic Acids Res 26: 4205–4213

Bannister AJ, Kouzarides T (2011) Regulation of chromatin by histone modifications. Cell Res 21: 381–395

Cao JJ, Wang TK, Wang Q, Zheng XW, Huang L (2019) Functional Insights Into Protein Acetylation in the Hyperthermophilic Archaeon. Molecular & Cellular Proteomics 18: 1572–1587

Capella-Gutiérrez S, Silla-Martínez JM, Gabaldón T (2009) trimAl: a tool for automated alignment trimming in large-scale phylogenetic analyses. Bioinformatics 25: 1972–1973

Cheng R, Huang FT, Lu XL, Yan Y, Yu BB, Wang XL, Zhu B (2023) Prokaryotic Gabija complex senses and executes nucleotide depletion and DNA cleavage for antiviral defense. Cell Host Microbe 31

Cheng R, Huang FT, Wu H, Lu XL, Yan Y, Yu BB, Wang XLE, Zhu B (2021) A nucleotide-sensing endonuclease from the Gabija bacterial defense system. Nucleic Acids Res 49: 5216–5229

Counts JA, Willard DJ, Kelly RM (2021) Life in hot acid: a genome-based reassessment of the archaeal order. Environ Microbiol 23: 3568–3584

Deng L, Zhu HJ, Chen ZJ, Liang YX, She QX (2009) Unmarked gene deletion and host-vector system for the hyperthermophilic crenarchaeon Sulfolobus islandicus. Extremophiles 13: 735–746

Doron S, Melamed S, Ofir G, Leavitt A, Lopatina A, Keren M, Amitai G, Sorek R (2018) Systematic discovery of antiphage defense systems in the microbial pangenome. Science 359

Dot EW, Thomason LC, Chappie JS (2023) Everything OLD is new again: How structural, functional, and bioinformatic advances have redefined a neglected nuclease family. Mol Microbiol 120: 122–140

Fan Y, Sanyal S, Bruzzone R (2018) Breaking Bad: How Viruses Subvert the Cell Cycle. Front Cell Infect Mi 8

Gao LY, Altae-Tran H, Böhning F, Makarova KS, Segel M, Schmid-Burgk JL, Koob J, Wolf YI, Koonin EV, Zhang F (2020) Diverse enzymatic activities mediate antiviral immunity in prokaryotes. Science 369: 1077-+

Gomez-Raya-Vilanova MV, Teuliere J, Medvedeva S, Dai Y, Corel E, Lopez P, Lapointe FJ, Bhattacharya D, Haraoui LP, Turc E et al (2025) Transcriptional landscape of the cell cycle in a model thermoacidophilic archaeon reveals similarities to eukaryotes. Nat Commun 16: 5697

Guindon S, Dufayard JF, Lefort V, Anisimova M, Hordijk W, Gascuel O (2010) New Algorithms and Methods to Estimate Maximum-Likelihood Phylogenies: Assessing the Performance of PhyML 3.0. Syst Biol 59: 307–321

Huo YW, Kong LF, Zhang Y, Xiao M, Du K, Xu SYT, Yan XX, Ma J, Wei TT (2024) Structural and biochemical insights into the mechanism of the Gabija bacterial immunity system. Nat Commun 15

Imachi H, Nobu MK, Nakahara N, Morono Y, Ogawara M, Takaki Y, Takano Y, Uematsu K, Ikuta T, Ito M et al (2020) Isolation of an archaeon at the prokaryote-eukaryote interface. Nature 577: 519-+

Katoh K, Rozewicki J, Yamada KD (2019) MAFFT online service: multiple sequence alignment, interactive sequence choice and visualization. Brief Bioinform 20: 1160–1166

Kiro R, Molshanski-Mor S, Yosef I, Milam SL, Erickson HP, Qimron U (2013) Gene product 0.4 increases bacteriophage T7 competitiveness by inhibiting host cell division. P Natl Acad Sci USA 110: 19549–19554

Koonin EV, Makarova KS, Wolf YI (2017) Evolutionary Genomics of Defense Systems in Archaea and Bacteria. Annu Rev Microbiol 71: 233-+

Li J, Cheng R, Wang ZM, Yuan WL, Xiao J, Zhao XY, Du XR, Xia SY, Wang LR, Zhu B et al (2024) Structures and activation mechanism of the Gabija anti-phage system. Nature

Li XY, Lozano-Madueno C, Martinez-Alvarez L, Peng X (2023) A clade of RHH proteins ubiquitous in Sulfolobales and their viruses regulates cell cycle progression (vol 51, pg 1724, 2023). Nucleic Acids Res 51: 4100–4100

Li YJ, Pan SF, Zhang Y, Ren M, Feng MX, Peng N, Chen LM, Liang YX, She QX (2016) Harnessing Type I and Type III CRISPR-Cas systems for genome editing. Nucleic Acids Res 44

Lindahl G, Sironi G, Bialy H, Calendar R (1970) Bacteriophage Lambda - Abortive Infection of Bacteria Lysogenic for Phage-P2. P Natl Acad Sci USA 66: 587-&

Lindås AC, Bernander R (2013) The cell cycle of archaea. Nat Rev Microbiol 11: 627–638

Lindås AC, Karlsson EA, Lindgren MT, Ettema TJG, Bernander R (2008) A unique cell division machinery in the Archaea. P Natl Acad Sci USA 105: 18942–18946

Liu J, Lelek M, Yang Y, Salles A, Zimmer C, Shen Y, Krupovic M (2025a) A relay race of ESCRT-III paralogs drives cell division in a hyperthermophilic archaeon. mBio 16: e0099124

Liu JF, Cvirkaite-Krupovic V, Baquero DP, Yang YF, Zhang Q, Shen YL, Krupovic M (2021a) Virus-induced cell gigantism and asymmetric cell division in archaea. P Natl Acad Sci USA 118

Liu JF, Cvirkaite-Krupovic V, Commere PH, Yang YF, Zhou F, Forterre P, Shen YL, Krupovic M (2021b) Archaeal extracellular vesicles are produced in an ESCRT-dependent manner and promote gene transfer and nutrient cycling in extreme environments. Isme J 15: 2892–2905

Liu JF, Gao RX, Li CT, Ni JF, Yang ZJ, Zhang Q, Chen HN, Shen YL (2017) Functional assignment of multiple ESCRT-III homologs in cell division and budding in. Mol Microbiol 105: 540–553

Liu JF, Lelek M, Yang YF, Salles A, Zimmer C, Shen YL, Krupovic M (2025b) A relay race of ESCRT-III paralogs drives cell division in a hyperthermophilic archaeon. Mbio 16

Liu Y, Makarova KS, Huang WC, Wolf YI, Nikolskaya AN, Zhang XX, Cai MW, Zhang CJ, Xu W, Luo ZH et al (2021c) Expanded diversity of Asgard archaea and their relationships with eukaryotes. Nature 593: 553-+

Lundgren M, Bernander R (2005) Archaeal cell cycle progress. Curr Opin Microbiol 8: 662–668

Lundgren M, Malandrin L, Eriksson S, Huber H, Bernander R (2008) Cell cycle characteristics of: Unity among diversity. J Bacteriol 190: 5362–5367

Matthews HK, Bertoli C, de Bruin RAM (2022) Cell cycle control in cancer. Nat Rev Mol Cell Bio 23: 74–88

Mestre MR, González-Delgado A, Gutierrez-Rus LI, Martínez-Abarca F, Toro N (2020) Systematic prediction of genes functionally associated with bacterial retrons and classification of the encoded tripartite systems. Nucleic Acids Res 48: 12632–12647

Millman A, Bernheim A, Stokar-Avihail A, Fedorenko T, Voichek M, Leavitt A, Oppenheimer-Shaanan Y, Sorek R (2020) Bacterial Retrons Function In Anti-Phage Defense. Cell 183: 1551-+

Minh BQ, Schmidt HA, Chernomor O, Schrempf D, Woodhams MD, von Haeseler A, Lanfear R (2020) IQ-TREE 2: New Models and Efficient Methods for Phylogenetic Inference in the Genomic Era. Mol Biol Evol 37: 1530–1534

Monroe N, Han H, Gonciarz MD, Eckert DM, Karren MA, Whitby FG, Sundquist WI, Hill CP (2014) The Oligomeric State of the Active Vps4 AAA ATPase. J Mol Biol 426: 510–525

Oh H, Koo J, An SY, Hong SH, Suh JY, Bae E (2023) Structural and functional investigation of GajB protein in Gabija anti-phage defense. Nucleic Acids Res 51: 11941–11951

Peng N, Deng L, Mei YX, Jiang DQ, Hu YM, Awayez M, Liang YX, She QX (2012) A Synthetic Arabinose-Inducible Promoter Confers High Levels of Recombinant Protein Expression in Hyperthermophilic Archaeon. Appl Environ Microb 78: 5630–5637

Peng N, Han WY, Li YJ, Liang YX, She QX (2017) Genetic technologies for extremely thermophilic microorganisms of, the only genetically tractable genus of crenarchaea. Sci China Life Sci 60: 370–385

Peng WF, Feng MX, Feng X, Liang YX, She QX (2015) An archaeal CRISPR type III-B system exhibiting distinctive RNA targeting features and mediating dual RNA and DNA interference. Nucleic Acids Res 43: 406–417

Risa GT, Hurtig F, Bray S, Hafner AE, Harker-Kirschneck L, Faull P, Davis C, Papatziamou D, Mutavchiev DR, Fan C et al (2020) The proteasome controls ESCRT-III-mediated cell division in an archaeon. Science 369: 642-+

Rocha EPC, Bikard D (2022) Microbial defenses against mobile genetic elements and viruses: Who defends whom from what? Plos Biol 20

Rousset F, Depardieu F, Miele S, Dowding J, Laval AL, Lieberman E, Garry D, Rocha EPC, Bernheim A, Bikard D (2022) Phages and their satellites encode hotspots of antiviral systems. Cell Host Microbe 30: 740-+

Samson RY, Obita T, Freund SM, Williams RL, Bell SD (2008) A Role for the ESCRT System in Cell Division in Archaea. Science 322: 1710–1713

Schiltz CJ, Adams MC, Chappie JS (2020) The full-length structure of OLD defines the ATP hydrolysis properties and catalytic mechanism of Class 1 OLD family nucleases. Nucleic Acids Res 48: 2762–2776

Schiltz CJ, Lee A, Partlow EA, Hosford CJ, Chappie JS (2019) Structural characterization of Class 2 OLD family nucleases supports a two-metal catalysis mechanism for cleavage. Nucleic Acids Res 47: 9448–9463

Spang A, Caceres EF, Ettema TJG (2017) MICROBIAL GENOMICS Genomic exploration of the diversity, ecology, and evolution of the archaeal domain of life. Science 357

Stewart CR, Deery WJ, Egan ESK, Myles B, Petti AA (2013) The product of SPO1 gene 56 inhibits host cell division during infection of by bacteriophage SPO1. Virology 447: 249–253

Takemata N, Samson RY, Bell SD (2019) Physical and Functional Compartmentalization of Archaeal Chromosomes. Cell 179: 165-+

Vorontsov EA, Rensen E, Prangishvili D, Krupovic M, Chamot-Rooke J (2016) Abundant Lysine Methylation and N-Terminal Acetylation in Revealed by Bottom-Up and Top-Down Proteomics. Mol Cell Proteomics 15: 3388–3404

Wei J, Li Y (2023) CRISPR-based gene editing technology and its application in microbial engineering. Eng Microbiol 3: 100101

Wolffe AP, Hayes JJ (1999) Chromatin disruption and modification. Nucleic Acids Res 27: 711–720

Xia YZ, Li K, Li JJ, Wang TQ, Gu LC, Xun LY (2019) T5 exonuclease-dependent assembly offers a low-cost method for efficient cloning and site-directed mutagenesis. Nucleic Acids Res 47

Yang W (2011) Nucleases: diversity of structure, function and mechanism. Q Rev Biophys 44: 1–93

Yang XY, Shen ZF, Xie JL, Greenwald J, Marathe I, Lin QP, Xie WJ, Wysocki VH, Fu TM (2024) Molecular basis of Gabija anti-phage supramolecular assemblies. Nat Struct Mol Biol

Yang Y, Liu J, Fu X, Zhou F, Zhang S, Zhang X, Huang Q, Krupovic M, She Q, Ni J et al (2023a) A novel RHH family transcription factor aCcr1 and its viral homologs dictate cell cycle progression in archaea. Nucleic Acids Res 51: 1707–1723

Yang YF, Liu JF, Fu XF, Zhou F, Zhang S, Zhang XM, Huang QH, Krupovic M, She QX, Ni JF et al (2023b) A novel RHH family transcription factor aCcr1 and its viral homologs dictate cell cycle progression in archaea. Nucleic Acids Res 51: 1707–1723

Zink IA, Pfeifer K, Wimmer E, Sleytr UB, Schuster B, Schleper C (2019) CRISPR-mediated gene silencing reveals involvement of the archaeal S-layer in cell division and virus infection. Nat Commun 10

