## Appendix Figures and Tables.docx for "A new class of OLD family ATPases Cran1 functions in cell cycle progression in an archaeon"

Content：

Appendix Figure S1. The Cran1 is highly conserved in Sulfolobales.

Appendix Figure S2. Gene synteny analysis of Cran1 proteins in representative species.

Appendix Figure S3. aCcr1 binds to the promoter *cran1*.

Appendix Figure S4. The growth of the *in situ* Flag-tagged Cran1 strain is not affected.

Appendix Figure S5. Phenotypic analysis of SiRe_0087 knockout strains and Cran1 and SiRe_0087 co-expression strains.

Appendix Table S1. Plasmids and strains used in the current study.

Appendix Table S2. Oligonucleotides used as primers in the current study.

Appendix Table S3. List of the sequences used for the phylogeny analysis in this study.(Attached Excel file)

Appendix Table S4. Mass spectrometry data of putative co-purified proteins with in situ Flag-tagged Cran1.(Attached Excel file)

### Appendix Figures

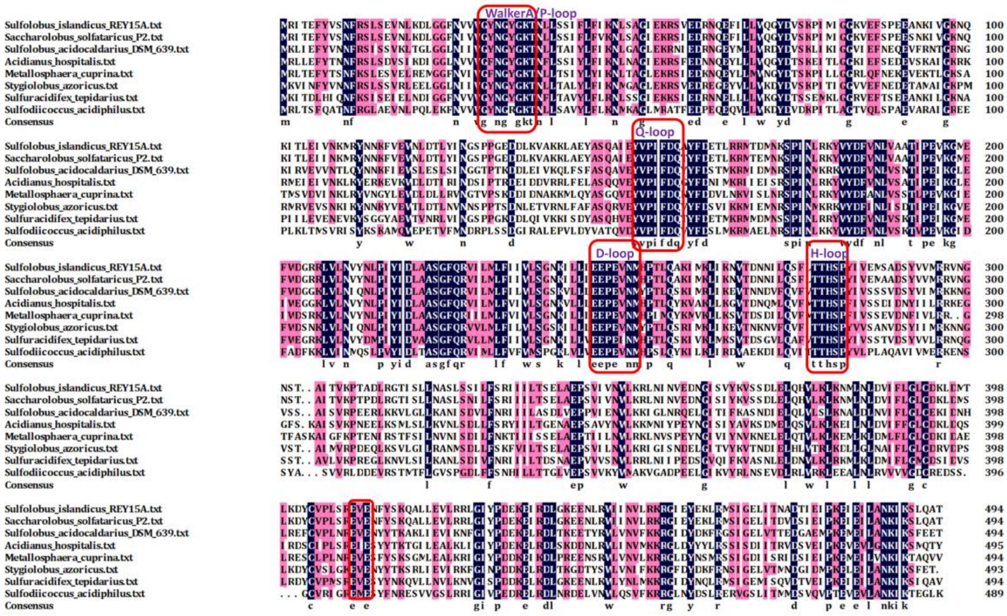


**Appendix Figure S1. The Cran1 is highly conserved in Sulfolobales.** Multiple sequence alignment of Cran1 homologs from Sulfolobales by DNAMAN software. The names of the species and strain where the homolog sequences were retrieved from the geneomes are indicated on the left. The protein sequences including Cran1 from S. islandicus REY15A (SiRe_0086), Sulfolobus acidocaldarius DSM639 (Saci_0157), Saccharolobus solfataricus,Acidianus, Metallosphaera cuprina, Sulfuracidifex tepidarius , Stygiolobus azoricus, Thermoprotei archaeon were retrieved from NCBI database after BLAST/PSI-BLAST analysis using SiRe_0086 as a query sequence.The red boxes are labelled with predicted conserved ATPase active sites, including P-loop,Q-loop,D-loop,H-loop.

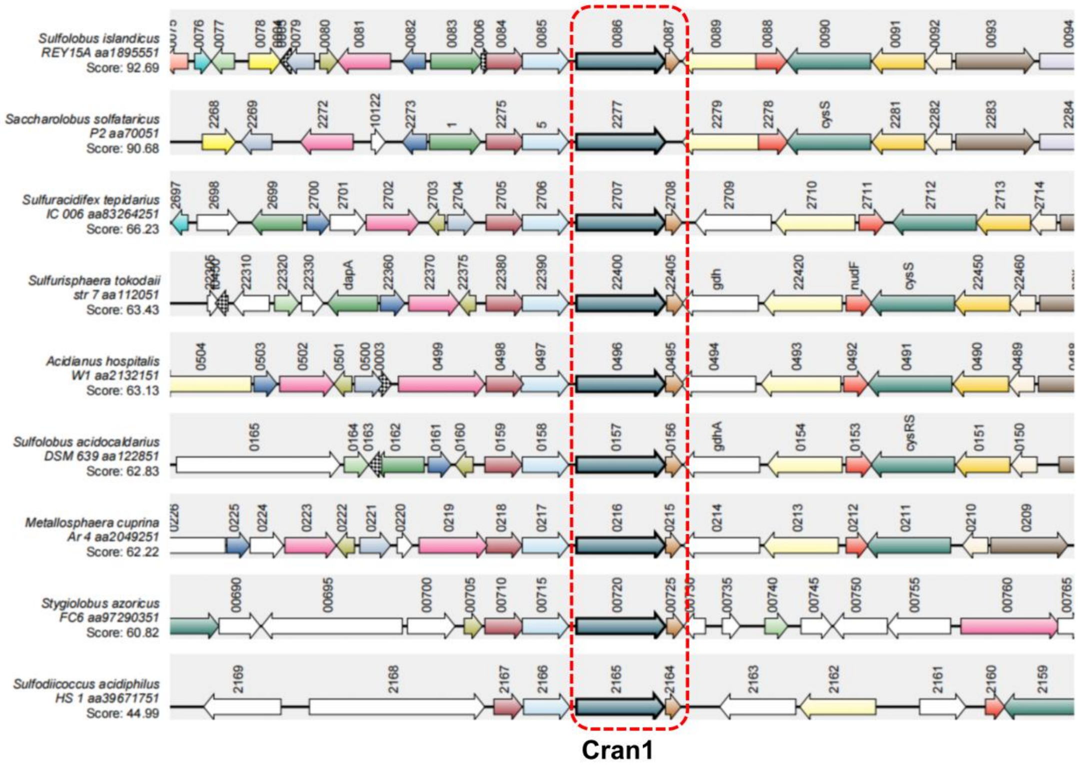


**Appendix Figure S2. Gene synteny analysis of Cran1 proteins in representative species.** The amino acid sequence of Cran1 (SiRe_0086) from *S. islandicus* REY15A was used as the query for the analysis by the SyntTax (https://archaea.i2bc.paris-saclay.fr/SyntTax/). The homologous genes corresponding to the query proteins are indicated with bold frames. The scores below the microbial names on the left indicate the normalized blast score between the query sequence and the orthologues in the matched chromosome.

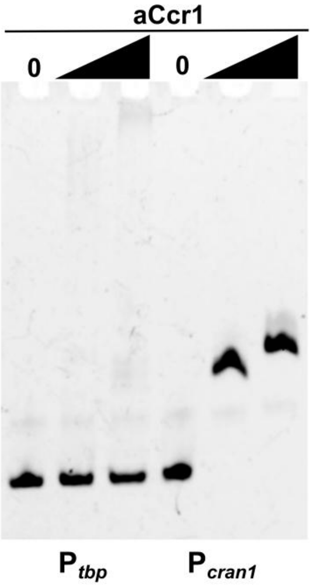


**Appendix Figure S3. aCcr1 binds to the promoter of *cran1*.** EMSA of aCcr1 binding to the promoter of *cran1* and *tbp* was shown. The 5' FAM-labelled and the corresponding complementary nucleotide sequences are listed in Appendix Table S2. The reaction was performed at 37°C for 30 min and analysed on a 10% native PAGE (see 'Materials and Methods'). Each reaction contained 2 nM of the FAM-labelled probe and 0, 0.5 and 1 μM aCcr1 protein.

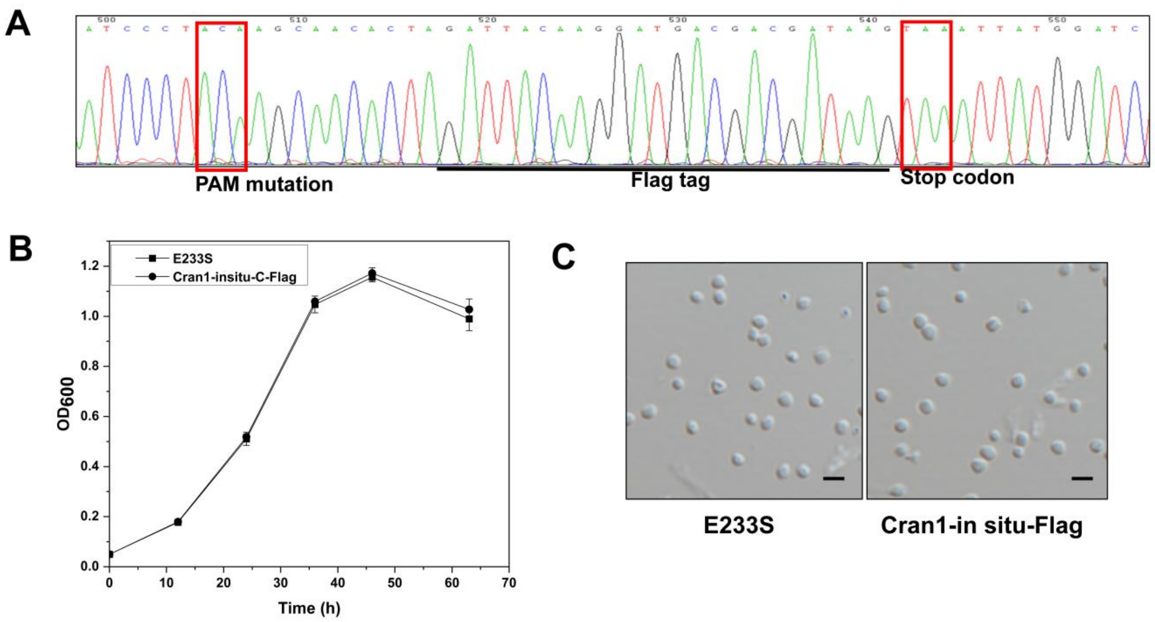


**Appendix Figure S4. The growth of the *in situ* Flag-tagged Cran1 strain is not affected.** (**A**) Verification of the *in situ* Flag-tagged strains by sequencing. The red boxes indicate the mutated PAM sequence and the stop codon, respectively. Comparison of the growth (**B**) and cell morphology (**C**) of the *in situ* Flag-tagged strain and the control E233S.

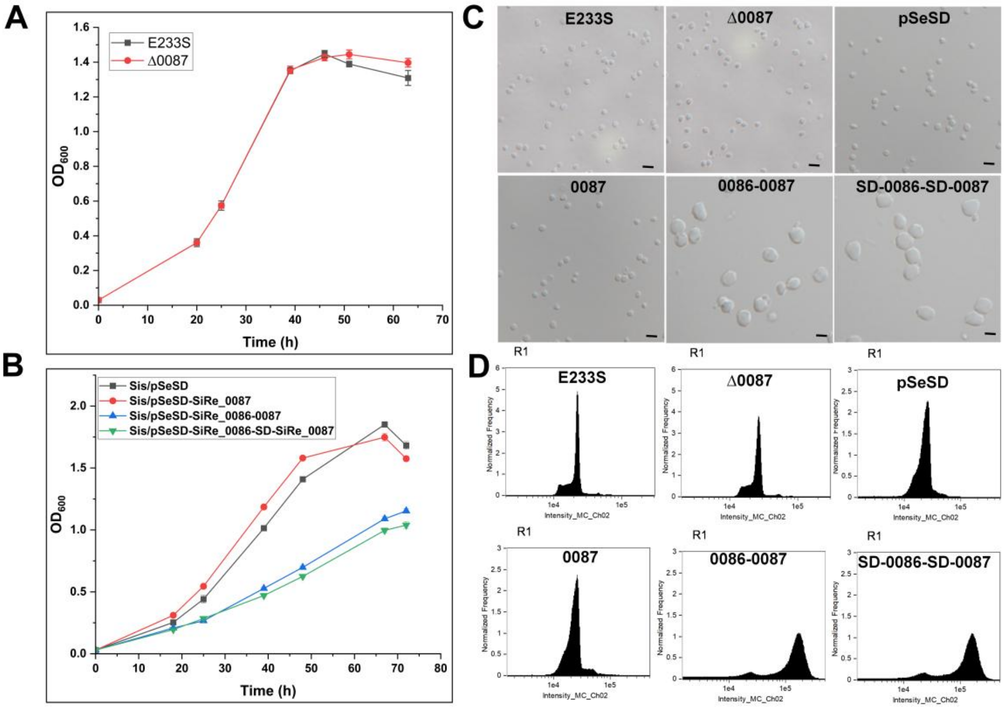


**Appendix Figure S5. Phenotypic analysis of SiRe_0087 knockout strains and Cran1 and SiRe_0087 co-expression strains.** (**A**) Growth curves of SiRe_0087 knockout strain and wild-type strain E233S were determined in MTSVU medium. Initial $OD_{600}$=0.03 (**B**) Overexpression of SiRe_0087 alone, co-expression of Cran1 and SiRe_0087 (sharing an arabinose promoter as well as using the arabinose promoter separately) strains were determined by growth curves in MTAV medium with an initial $OD_{600}$=0.03. (**C**) Observation of cellular morphology of the strains in A and B at 24h of growth and flow cytometry analysis (**D**) Scale bars: 2 μm.

### Appendix Tables

**Appendix Table S1.** Plasmids and strains used in the current study.

| **Strain/Plasmid** | **Description** | **References or sources** |
|---|---|---|
| pSeSD | A Sulfolobus-E. coli shuttle vector carrying an expression cassette controlled under a synthetic strong promoter ParaS-SD | (1) |
| pGE-Cran1-KO | The genome-editing plasmid for *cran1* knockout | This study |
| pGE-SiRe_0087-KO | The genome-editing plasmid for *sire_0087* knockout | This study |
| Sis/pGE-Cran1-insitu -Flag | E233S harboring the genome-editing plasmid for *cran1* add a C-terminal flag tag | This study |
| Sis/pGE-Cran1-insitu-His | E233S harboring the genome-editing plasmid for *cran1* add a C-terminal His tag | This study |
| Cran1-insitu-flag/pGE-Cran1-KD-4S | Cran1-insitu-flag strains harboring the genome-editing plasmid for *cran1* knockdown | This study |
| Sis/pSeSD | E233S harboring pSeSD plasmid | This study |
| Sis/pSeSD-Cran1-C-His | E233S harboring pSeSD-Cran1-C-His | This study |
| Sis/pSeSD-Cran1-no tag | E233S harboring pSeSD-Cran1-no tag | This study |
| Sis/pSeSD-SiRe_0087-no tag | E233S harboring pSeSD-SiRe_0087-no tag | This study |
| Sis/pSeSD-SiRe_0086-SiRe_0087-no tag | E233S harboring SiRe_0086 and SiRe_0087 co-expression plasmids | This study |
| Sis/pSeSD-Cran1-K36A-C-His | E233S harboring pSeSD-Cran1-K36A-C-His | This study |
| Sis/pSeSD-Cran1-T37A-C-His | E233S harboring pSeSD-Cran1-T37A-C-His | This study |
| Sis/pSeSD-Cran1-F156A-C-His | E233S harboring pSeSD-Cran1-F156A-C-His | This study |
| Sis/pSeSD-Cran1-K36A-C-His | E233S harboring pSeSD-Cran1-K36A-C-His | This study |

| | | |
|---|---|---|
| Sis/pSeSD-Cran1-T37A-C-His | E233S harboring pSeSD-Cran1-T37A-C-His | This study |
| Sis/pSeSD-Cran1-F156A-C-His | E233S harboring pSeSD-Cran1-F156A-C-His | This study |
| Sis/pSeSD-Cran1-D157A-C-His | E233S harboring pSeSD-Cran1-D157A-C-His | This study |
| Sis/pSeSD-Cran1-Q158A-C-His | E233S harboring pSeSD-Cran1-Q158A-C-His | This study |
| Sis/pSeSD-Cran1-P246A-C-His | E233S harboring pSeSD-Cran1-P246A-C-His | This study |
| Sis/pSeSD-Cran1-E247A-C-His | E233S harboring pSeSD-Cran1-E247A-C-His | This study |
| Sis/pSeSD-Cran1-P248A-C-His | E233S harboring pSeSD-Cran1-P248A-C-His | This study |
| Sis/pSeSD-Cran1-E249A-C-His | E233S harboring pSeSD-Cran1-E249A-C-His | This study |
| Sis/pSeSD-Cran1-H280A-C-His | E233S harboring pSeSD-Cran1-H280A-C-His | This study |
| Sis/pSeSD-Cran1-S281A-C-His | E233S harboring pSeSD-Cran1-S281A-C-His | This study |
| Sis/pSeSD-Cran1-T37A/H280A/S281A-C-His | E233S harboring pSeSD-Cran1-T37A/H280A/S281A-C-His | This study |
| Sis/pSeSD-Cran1-E338A/L391A/D393A-C-His | E233S harboring pSeSD-Cran1-E338A/L391A/D393A-C-His | This study |
| Sis/pSeSD-Cran1-E341A/E409A/E411A-C-His | E233S harboring pSeSD-Cran1-E341A/E409A/E411A-C-His | This study |
| Sis/pSeSD-Cran1-L391A/D393A/E409A/E411A-C-His | E233S harboring pSeSD-Cran1-L391A/D393A/E409A/E411A-C-His | This study |

| | | |
|---|---|---|
| Sis/pSeSD-Saci_OLD | E233S harboring pSeSD-Saci_OLD strain | This study |
| *cran1-Flag*/pGE | Cran1 genome in situ Flag-tagged strain carrying pGE empty vector | This study |
| *cran1-Flag*/ pGE-KD-4spacers | Cran1 genome in situ Flag-tagged strain carrying pGE-Cran1 knockdown vector | This study |
| *E. coli* /pET22b-Cran1-C-His | Expression of WT-Cran1 having His-tag at the C-terminal | This study |
| *E. coli* /pET22b-Cran1-K36A-C-His | Expression of Cran1-K36A having His-tag at the C-terminal | This study |
| *E. coli* /pET22b-Cran1-T37A-C-His | Expression of Cran1-T37A having His-tag at the C-terminal | This study |
| *E. coli* /pET22b-Cran1-F156A-C-His | Expression of Cran1-F156A having His-tag at the C-terminal | This study |
| *E. coli* /pET22b-Cran1-D157A-C-His | Expression of Cran1-D157A having His-tag at the C-terminal | This study |
| *E. coli* /pET22b-Cran1-Q158A-C-His | Expression of Cran1-Q158A having His-tag at the C-terminal | This study |
| *E. coli* /pET22b-Cran1-P248A-C-His | Expression of Cran1-P248A having His-tag at the C-terminal | This study |
| *E. coli* /pET22b-Cran1-E249A-C-His | Expression of Cran1-E249A having His-tag at the C-terminal | This study |
| *E. coli* /pET22b-Cran1-H280A-C-His | Expression of Cran1-H280A having His-tag at the C-terminal | This study |
| *E. coli* /pET22b-Cran1-S281A-C-His | Expression of Cran1-S281A having His-tag at the C-terminal | This study |
| *E. coli* /pET22b-Cran1-T37A/H280A/S281A-C-His | Expression of Cran1-T37A/H280A/S281A having His-tag at the C-terminal | This study |

| | | |
|---|---|---|
| *E. coli* /pET22b-Cran1-E338A/L391A /D393A-C-His | Expression of Cran1-E338A/L391A/D393A having His-tag at the C-terminal | This study |
| *E. coli* /pET22b-Cran1-E341A/E409A /E411A-C-His | Expression of -Cran1-E341A/E409A/E411A having His-tag at the C-terminal | This study |
| *E. coli* /pET22b--Cran1-L391A/D393 A/E409A/E411A-C-His | Expression of -Cran1-L391A/D393A/E409A/E411A having His-tag at the C-terminal | This study |
| *MG1655*/PET22b | *MG1655* strain carrying the PET22b empty vector | This study |
| *MG1655*/PET22b-SiRe_0086 | *MG1655* strain carrying the PET22b-SiRe_0086 vector | This study |
| *MG1655*/PET22b-SiRe_0086-0087 | *MG1655* strain carrying the PET22b-SiRe_0086-0087 vector | This study |

**Appendix Table S2.** Oligonucleotides used as primers in the current study.

| Primer | Sequence [a, b] (5´-3´) |
|---|---|
| Cran1-KO-spacer-F | AAGACTCTTCAAGCTAAAATCATGAAATTAATTAAGAACTGGA |
| Cran1-KO-spacer-R | AGCTCCAGTTCTTAATTAATTTCATGATTTTAGCTTGAAGAGT |
| Cran1-L-arm-F-sphI | AAGTACAATTGTGCTGCATGCGACAAAGACGGAAAGGCT |
| Cran1-L-arm-R | AACATTTATATTCCATCTTCTATTCCGGCTGACAAA |
| Cran1-R-arm-F | TTTGTCAGCCGGAATAGA AGATGGATTATAAATGTT |
| Cran1-R-arm-R-XhoI | TTAACATATTGGATGCTCGAGTTTCATTTACCATCTCTA |
| Cran1Flanking-F | GCTTACGTTACCGGTAGTCCAAAAC |
| Cran1Flanking-R | CTTCATAGCATACCAGTACGTTCG |
| Cran1-in situ-Sp-F | AAGAGCAACACTATAAATTATGGATCTTAGTATCTTATTCGAT |
| Cran1-in situ-Sp-R | AGCATCGAATAAGATACTAAGATCCATAATTTATAGTGTTGCT |
| Cran1insitu-L-arm-F | AAGTACAATTGTGCTGCATGCTTAGTAGGATTATAATACTG |
| Cran1insitu-L-arm-R | CTTATCGTCGTCATCCTTGTAATCTAGTGTTGCTTGTAGGGATT |
| Cran1insitu-R-arm-F | GATTACAAGGATGACGACGATAAGTAAATTATGGATCTTAGTAT |
| Cran1insitu-R-arm-R | TTAACATATTGGATGCTCGAGAAGAAGTAGTTGAGATGATT |
| Cran1insitu-L-His-R | ATGGTGATGGTGATGATGATGGTGATGGTGATGATGTAGTGTTGC TTGTAGGGATT |
| Cran1insitu-R-His-F | CATCATCACCATCACCATCATCATCACCATCACCATTAAATTATGG ATCTTAGTAT |
| Flag-F | GATTACAAGGATGACGACGATAAG |
| His-F | ATGGTGATGGTGATGATG |
| Cran1-NdeI-F | GAATGAGGTGAAGCTCATATGAGGATTACTGAATTTTAC |
| Cran1-SalI-R | GGCCGCTTGATCAGCGTCGACTAGTGTTGCTTGGAGGG |
| Cran1-ST-salI-R | GGCCGCTTGATCAGCGTCGACTTATAGTGTTGCTTGGAGGGATT |
| Cran1K36A-F | AACGGATATGGA**GCA**ACTAATCTACTT |
| Cran1K36A-R | AAGTAGATTAGT**TGC**TCCATATCCGTT |
| Cran1T37A-F | AACGGATATGGAAAA**GCT**AATCTACTT |

| | |
|---|---|
| Cran1T37A-R | AAGTAGATT**AGC**TTTTCCATATCCGTT |
| Cran1F156A-F | TGTACCTATC**GCC**GACCAAGCGTATTTTG |
| Cran1F156A-R | CAAAATACGCTTGGTC**GGC**GATAGGTACA |
| Cran1D157A-F | TGTACCTATCTTC**GCC**CAAGCGTATTTTG |
| Cran1D157A-R | CAAAATACGCTTG**GGC**GAAGATAGGTACA |
| Cran1Q158A-F | TGTACCTATCTTCGAC**GCA**GCGTATTTTG |
| Cran1Q158A-R | CAAAATACGC**TGC**GTCGAAGATAGGTACA |
| Cran1P246A-F | AAGATATTACTAATT**GCA**GAACCAGAGGTGAA |
| Cran1P246A-R | TTCACCTCTGGTTC**TGC**AATTAGTAATATCTT |
| Cran1E247A-F | AAGATATTACTAATTGAA**GCA**CCAGAGGTGAA |
| Cran1E247A-R | TTCACCTCTGG**TGC**TTCAATTAGTAATATCTT |
| Cran1P248A-F | AATTGAAGAA**GCA**GAGGTGAATATGC |
| Cran1P248A-R | GCATATTCACCTC**TGC**TTCTTCAATT |
| Cran1E249A-F | AATTGAAGAACCA**GCG**GTGAATATGC |
| Cran1E249A-R | GCATATTCAC**CGC**TGGTTCTTCAATT |
| Cran1H280A-F | CCTTACAACA**GCT**TCCCCATACATCG |
| Cran1H280A-R | CGATGTATGGGGA**AGC**TGTTGTAAGG |
| Cran1S281A-F | CCTTACAACACAT**GCC**CCATACATCG |
| Cran1S281A-R | CGATGTATGG**GGC**ATGTGTTGTAAGG |
| Cran1H280AS281A-F | CCTTACAACA**GCTGCC**CCATACATCG |
| Cran1H280AS281A-R | CGATGTATGG**GGCAGC**TGTTGTAAGG |
| Cran1-E338A-F | ATACTGACAAGT**GCG**TTAGCAGAA |
| Cran1-E338A-R | TTCTGCTAA**CGC**ACTTGTCAGTAT |
| Cran1-L391A-D393A-F | CGTGATATTCCTTGGA**GCA**TGT**GCC**AAGCTTGATATGACCTTAAAGGAT |
| Cran1-L391A-D393A-R | ATCCTTTAAGGTCATATCAAGCTT**GGC**ACA**TGC**TCCAAGGAATATCACG |
| Cran1-E341A-F | CAAGTGAGTTAGCA**GCA**CCCTCTGTAATAGTGAAT |
| Cran1-E341A-R | ATTCACTATTACAGAGGG**TGC**TGCTAACTCACTTG |

| | |
|---|---|
| Cran1-E409A-E411A-F | GCGTTCCTTTAAGTCGT**GCG**GTA**GCG**AATTTCTATAGTAAACAAG |
| Cran1-E409A-E411A-R | CTTGTTTACTATAGAAATT**CGC**TAC**CGC**ACGACTTAAAGGAACGC |
| Sac_0157-NdeI-F | GAATGAGGTGAAGCTCATATGAAGCTTATTGAGTTTTACGT |
| Sac_0157-SalI-R | GGCCGCTTGATCAGCGTCGACTATAGTTTCTTCGAAAGATT |
| Cran1KD-F1 | GCTAATCTACTATAGAATTGAAAGTAGATTAGTTTTTCCATATCCGTTATAGCCAACGA |
| Cran1KD-F2 | CTACGGCTAATCTACTATAGAATTGAAAGTAGATTAGTTTTTCCATATCCGTTATAGCC |
| Cran1KD-R1 | CTAATCTACTTTCAATTCTATAGTAGATTAGCCGTAGTCGTTGGCTATAACGGATATGG |
| Cran1KD-R2 | CTTTCAATTCTATAGTAGATTAGCCGTAGTCGTTGGCTATAACGGATATGGAAAAA |
| 0087-SalI-ST-R | GGCCGCTTGATCAGCGTCGACCTACAAGATGTAAAATATCA |
| 0087-NdeI-F | GAATGAGGTGAAGCTCATATGATGGATCTTAGTATCTTATT |
| FAM-$P_{Cran1}$-F | AGATAATACTCCGTTTTTCTCTAACAATAGATTAACGTTAAAGAAGCCTGCTAACGATTATATTTTATTCTGCTAAGTATTACCGTAGAGTGATCTGTA |
| $P_{Cran1}$-R | TACAGATCACTCTACGGTAATACTTAGCAGAATAAAATATAATCGTTAGCAGGCTTCTTTAACGTTAATCTATTGTTAGAGAAAAACGGAGTATTATCT |
| 0087KOspacer-F | AAGTCTGATAGGGATCAAAAAACACATGTTGGTGGGGTAATTT |
| 0087KOspacer-R | AGCAAATTACCCCACCAACATGTGTTTTTTGATCCCTATCAGA |
| 0087KOL-arm-sphI-F | AAGTACAATTGTGCTGCATGCGGATTATAATACTGACAAGT |
| 0087KOL-arm-R | AATTTTTTATCATAATCACGAATTTATAGTGTTGCTTGGA |
| 0087KOR-arm-F | TCCAAGCAACACTATAAATTCGTGATTATGATAAAAAATT |

| | |
|---|---|
| 0087KOR-arm-xhoI-R | TTAACATATTGGATGCTCGAGCGGCTTCATTATTGGGAGAT |
| 0087KO-Flanking-F | GTCTTTCCTTACAACACATT |
| 0087KO-Flanking-R | GGTTATTTGCAAATAAGTTA |

[a] The underlined denote sites of restriction enzymes. [b] The mutated codons are indicated in boldface.

1. Peng, N., Deng, L., Mei, Y.X., Jiang, D.Q., Hu, Y.M., Awayez, M., Liang, Y.X. and She, Q.X. (2012) A Synthetic Arabinose-Inducible Promoter Confers High Levels of Recombinant Protein Expression in Hyperthermophilic Archaeon. *Appl Environ Microb*, **78**, 5630-5637.

**Appendix Table S3.** List of the sequences used for the phylogeny analysis in this study.

**Appendix Table S4.** Mass spectrometry data of putative co-purified proteins with in situ Flag-tagged Cran1
